# Organization and dynamics of the cortical complexes controlling insulin secretion in β-cells

**DOI:** 10.1101/2021.08.11.455909

**Authors:** Ivar Noordstra, Cyntha M. van den Berg, Fransje W. J. Boot, Eugene A. Katrukha, Ka Lou Yu, Roderick P. Tas, Sybren Portegies, Bastiaan J. Viergever, Esther de Graaff, Casper C. Hoogenraad, Eelco J. P. de Koning, Françoise Carlotti, Lukas C. Kapitein, Anna Akhmanova

## Abstract

Insulin secretion in pancreatic β-cells is regulated by cortical complexes that are enriched at the sites of adhesion to extracellular matrix facing the vasculature. Many components of these complexes, including Bassoon, RIM, ELKS and liprins, are shared with neuronal synapses. Here, we show that insulin secretion sites also contain non-neuronal proteins LL5β and KANK1, which in migrating cells organize exocytotic machinery in the vicinity of integrin-based adhesions. Depletion of LL5β or focal adhesion disassembly triggered by myosin II inhibition perturbed the clustering of secretory complexes and attenuated the first wave of insulin release. While previous analyses in vitro and in neurons suggested that secretory machinery might assemble through liquid-liquid phase separation, analysis of endogenously labeled ELKS in pancreatic islets indicated that its dynamics is inconsistent with such a scenario. Instead, fluorescence recovery after photobleaching and single molecule imaging showed that ELKS turnover is driven by binding and unbinding to low-mobility scaffolds. Both the scaffold movements and ELKS exchange were stimulated by glucose treatment. Our findings help to explain how integrin-based adhesions control spatial organization of glucose-stimulated insulin release.

## Introduction

Pancreatic β-cells respond to elevated glucose levels in the bloodstream by the activation of exocytotic machinery and the release of insulin, which subsequently stimulates glucose uptake and conversion in different tissues (Meglasson et al., 1986). Defects in insulin secretion lead to diabetes type II, a major worldwide health problem, which affects increasingly large numbers of patients.

Insulin exocytosis is regulated by specialized cortical complexes, which contain Bassoon, Piccolo, RIM, ELKS and liprins (Gan et al., 2017, Low et al., 2014, Ohara-Imaizumi et al., 2019b), reviewed in (Ohara-Imaizumi et al., 2019a). These proteins are also present in neurons, where they form the cytomatrix at the active zone (CAZ), a dense protein meshwork that spatially organizes the coupling of calcium influx to neurotransmitter release (Gundelfinger et al., 2012, Sudhof, 2012). However, unlike neuronal synapses, the secretion sites in pancreatic β-cells are enriched at the interface with vasculature, where β-cells make contacts with the extracellular matrix (Cottle et al., 2021, Gan et al., 2018, Gan et al., 2017, Low et al., 2014). Interestingly, ELKS and liprins, but not the other CAZ proteins, are also part of the cortical complexes regulating constitutive, calcium-independent secretion in non-neuronal cells, such as fibroblasts, keratinocytes and cancer cells (Grigoriev et al., 2007, Lansbergen et al., 2006, Stehbens et al., 2014, van der Vaart et al., 2013). In such cells, the localization of secretory machinery is controlled by integrin-based cell adhesion, and secretory complexes are concentrated at leading cell edges and around focal adhesions (Fourriere et al., 2019, Grigoriev et al., 2007, Stehbens et al., 2014). Integrin dependent adhesion, integrin activation and focal adhesion signaling play an important role in insulin secretion ((Bosco et al., 2000, Cai et al., 2012, Gan et al., 2018, Kaido et al., 2006, Krishnamurthy et al., 2008, Parnaud et al., 2009, Parnaud et al., 2006, Riopel et al., 2011, Riopel et al., 2013, Rondas et al., 2011, Rondas et al., 2012), reviewed in (Arous et al., 2015, Arous et al., 2017, Kalwat et al., 2013)). Specifically, glucose stimulation of β-cells leads to the myosin IIA-dependent remodeling of F-actin and the activation of focal adhesion kinase (FAK), paxillin and ERK, ultimately resulting in focal adhesion enlargement (Arous & Halban, 2015, Arous et al., 2013, Gan et al., 2018, Rondas et al., 2011, Rondas et al., 2012, Wilson et al., 1999, Wilson et al., 2001). Although the importance of focal adhesion remodeling for insulin secretion has been confirmed *in vivo* (Cai et al., 2012), little is known about the relationship between focal adhesions and the proteins organizing insulin exocytosis. Furthermore, a flurry of recent studies suggested that in neurons, CAZ may be formed by liquid-liquid phase separation (LLPS) of the key players, such as liprins, ELKS and RIMs (Emperador-Melero et al., 2021, Liang et al., 2021, McDonald et al., 2020, Sala et al., 2019, Wu et al., 2019, Wu et al., 2021); reviewed in (Chen et al., 2020, Hayashi et al., 2021)). The idea that the exocytotic machinery in β-cells would form by LLPS is attractive, but it has not been tested.

Here, we explored the connections between the secretory machinery and focal adhesions in β-cells and investigated whether secretory complexes behave like liquid condensates. Previous work showed that two non-neuronal proteins, LL5β and KANK1, which interact with phosphatidylinositol (3,4,5)-trisphosphate and the integrin adhesion component talin, respectively, play a key role in organizing exocytotic complexes around focal adhesions in different cell types (Bouchet et al., 2016a, Grigoriev et al., 2007, Lansbergen et al., 2006, Stehbens et al., 2014, van der Vaart et al., 2013). We found that both proteins are part of exocytotic complexes and that LL5β is required for efficient clustering of secretory complex components in β-cells. We showed that such clustering occurs around focal adhesions and depends on the actomyosin contractility. Importantly, perturbation of the clustering of secretory complexes led to the inhibition of the first, rapid phase of insulin secretion, which is known to depend on the release of the pre-docked pool of insulin granules (Rorsman et al., 2003, Wang et al., 2009). To test whether the clustering of secretory complexes is driven by LLPS, we focused on the dynamics of ELKS (also known as ELKS1, ERC1, RAB6 Interacting Protein 2 and CAST2), because ELKS and is homologue CAST/ELKS2/ERC2 are multivalent proteins that can interact with multiple components of the secretory machinery, including liprin-α isoforms, RIM, Bassoon, LL5β and voltage-gated calcium channels (Kiyonaka et al., 2012, Ko et al., 2003, Lansbergen et al., 2006, Takao-Rikitsu et al., 2004, Wang et al., 2002). We used a newly generated GFP-ELKS mouse knock-in model to investigate in detail the dynamics of endogenously labelled secretory complexes in pancreatic islets. We confirmed the colocalization of neuronal and non-neuronal proteins at the insulin secretion sites in the vicinity of blood vessels in pancreatic islets, characterized the dynamics of ELKS-containing secretory complexes and demonstrated that their turnover is strongly regulated by glucose levels. We found no evidence that would support LLPS as the basis for formation of ELKS foci, because the majority of them contained only 2-4 ELKS dimers and showed all hallmarks of protein binding/unbinding to a low-mobility scaffold rather than liquid condensates. Our data support the view that formation secretory sites in pancreatic β-cells is driven by protein binding and unbinding to scaffolds clustered around focal adhesions.

## Results

### LL5β and KANK1 colocalize with CAZ components in pancreatic β-cells

Using insulin secreting INS-1E cells as a model system, we confirmed that the regions of cortical accumulation of insulin granules corresponded to the areas where CAZ markers, such as RIM1 or RIM2 (detected here with antibodies recognizing both proteins, which will be termed RIM from now on) and Bassoon, are enriched (Fig. 1A,B, Fig. S1A). Next, we tested whether the localization of LL5β and KANK1 overlaps with that of RIM, Bassoon, liprins and ELKS. Imaging of the cell cortex by Total Internal Reflection Fluorescence microscopy (TIRFM) showed that all these proteins displayed punctate staining patterns that were concentrated in the same membrane regions (Fig. 1C,D, Fig. S1B-D). These data confirm previous observations on the colocalization between LL5β, liprins, ELKS and KANK1 in different cell types (Astro et al., 2014, Bernadzki et al., 2014, Bouchet et al., 2016a, Lansbergen et al., 2006, van der Vaart et al., 2013, Yuan et al., 2015) and show that in insulin-secreting cells, these proteins also colocalize with CAZ components. We also observed that the LL5β-binding microtubule-stabilizing protein CLASP1 was enriched in cell regions containing LL5β accumulations (Fig. S1E, Fig. 1D). A higher-resolution analysis using Stimulated Emission Depletion (STED) microscopy showed that the puncta observed with different antibodies were closely apposed but did not coincide (Fig. 1E, Fig. S1F). Presence of ELKS, Bassoon, RIM and LL5β in the same complexes in INS-1E cells was also confirmed by immunoprecipitation of endogenous proteins with anti-ELKS antibodies (Fig. 1F). The spatial distribution and colocalization of the cortical protein complexes and insulin was confirmed in the human β-cell line EndoC-βH1 (Fig. 1G,H, Fig. S1G) and dispersed human pancreatic islets (Fig. 1I,J, Fig. S1H). Taken together, these experiments reveal a conserved hybrid protein complex in pancreatic β-cells that consists of both neuronal CAZ components and non-neuronal proteins that regulate cortical microtubule attachment and constitutive secretion in epithelia and fibroblasts.

**Figure 1.**
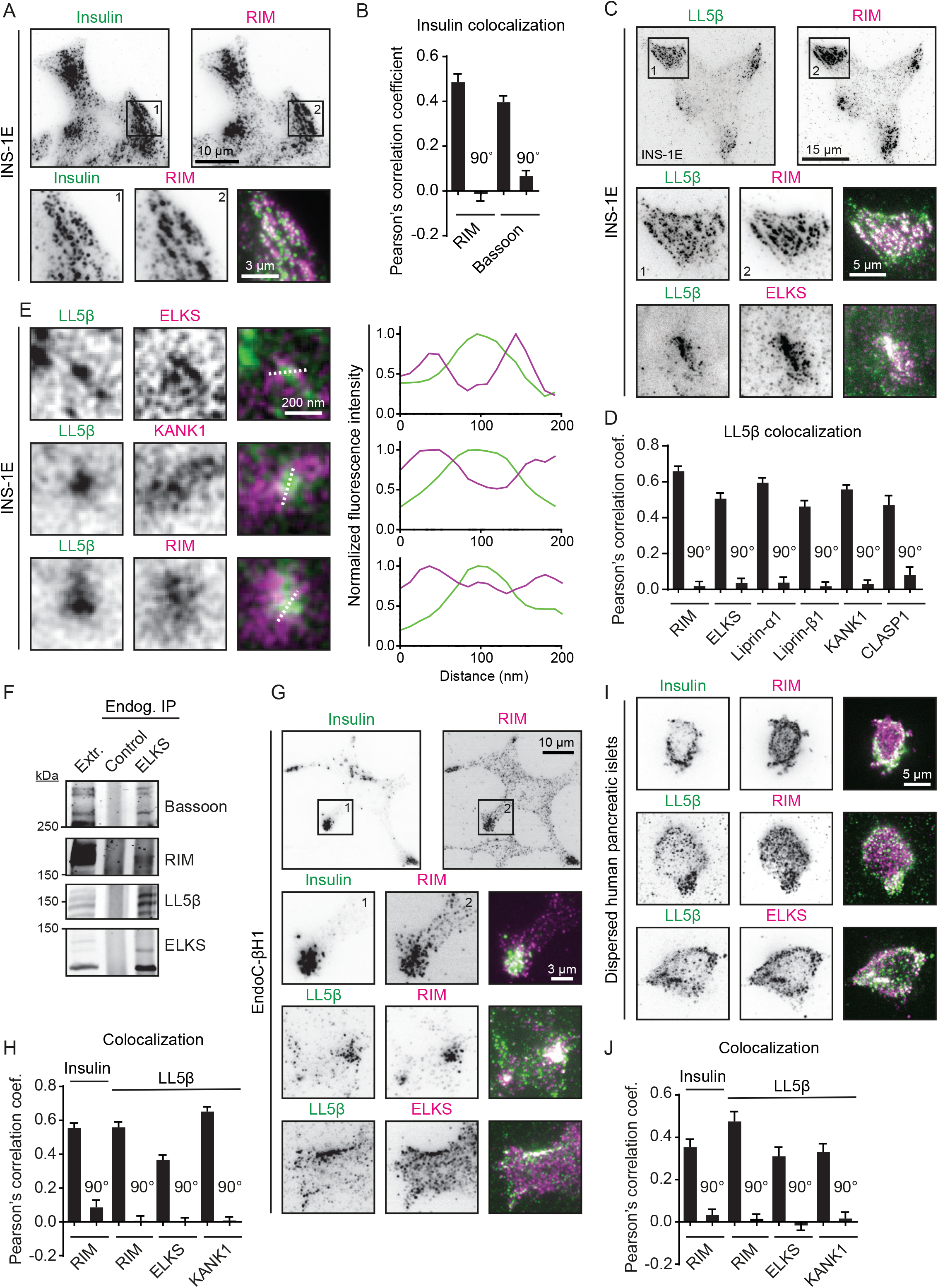
LL5β and KANK1 colocalize with CAZ components and insulin granules in pancreatic beta cell lines and dispersed human pancreatic islets. A. Staining for insulin (green) and RIM (RIM1 and RIM2) (magenta) in INS-1E cells imaged with total internal reflection fluorescence microscopy (TIRFM). B. Quantification of colocalization between insulin and indicated proteins in INS-1E cells using Pearson’s correlation coefficient between the two channels. For analysis, intracellular regions of interest (ROIs) of approximately 25 µm^2^ were used. For RIM, n=15; for Bassoon, n=21; 90° indicates a 90° rotation of one of the two analyzed channels before analysis; error bars, S.E.M. C. Staining for LL5β (green) and RIM and ELKS (magenta) in INS-1E cells imaged with TIRFM. D. Quantification of colocalization between LL5β and indicated proteins in INS-1E cells. Analysis and display as in (B). For all conditions, n=15-33. E. Stimulated Emission Depletion (STED) microscopy images of LL5β (green) and ELKS, KANK1 and RIM (magenta) in INS-1E cells. Intensity profiles along dotted lines are plotted in graphs. F. Co-immunoprecipitation assay from INS-1E cell extracts using antibodies against endogenous ELKS. Rabbit IgG was used as a control. G. Staining for Insulin or LL5β (green) and RIM and ELKS (magenta) in EndoC-βH1 cells imaged with TIRFM. H. Quantification of colocalization between insulin or LL5β and indicated proteins in EndoC-βH1 cells using Pearson’s correlation coefficient between the two channels. Analysis and display as in (B). For all conditions, n=27-30. I. Staining for Insulin or LL5β (green) and RIM and ELKS (magenta) in dispersed human pancreatic islets imaged with TIRFM. J. Quantification of colocalization between insulin or LL5β and indicated proteins in dispersed human pancreatic islets using Pearson’s correlation coefficient between the two channels. Analysis and display as in (B). For all conditions, n=14-22.

### LL5β is required for clustering of insulin secretion complexes and efficient insulin release

Our previous work showed that in non-neuronal cells, LL5β, liprins and KANK1 can independently localize to the plasma membrane, but require each other as well as ELKS for the formation of dense cortical clusters at the leading cell edges and around focal adhesions (Bouchet et al., 2016a, Lansbergen et al., 2006, van der Vaart et al., 2013). To test if the same is true for insulin-secreting cells, we depleted LL5β in INS-1E cells using two different siRNAs and observed a 30-40% reduction in the LL5β signal on Western blots (Fig. S2A,B). Immunofluorescence cell staining showed that LL5β-positive puncta were almost completely lost in ∼30-40% of the cells (Fig. 2A,B). Depletion of LL5β did not prevent membrane localization of RIM, but its clustering was reduced (Fig. 2A,C,D). Next, we examined whether the depletion of LL5β affected glucose-stimulated insulin secretion. In control cells, the cortex-associated insulin granule pool was strongly reduced due to the rapid insulin secretion within the first 5 minutes after glucose stimulation, and then gradually recovered, as described previously (Curry et al., 1968, Rorsman et al., 2000) (Fig. S2C,D). Due to insufficient compatibility of cell fixation procedures required to detect LL5β and insulin granules in INS-1E cells, it was not possible to stain them simultaneously. We therefore co-stained INS-1E cells for insulin and RIM, and analyzed the cells with dispersed RIM puncta, as we observed that such cells were strongly depleted of LL5β (Fig. 2A). In the absence of glucose stimulation, the density of insulin granules in the vicinity of the basal cortex was not affected, however their release in the vicinity of the basal cortex after 5 minutes of glucose stimulation was inhibited in LL5β-depleted cells (Fig. 2E,F). Importantly, LL5β knockdown compromised only the secretion of the cortex-associated insulin granule pool, but did not affect insulin vesicle numbers further away from the basal cortex (Fig. 2G, H). These data suggest that LL5β-dependent clustering of the components of exocytotic machinery has no effect on the formation and possibly also docking of insulin granules but affects their fusion and thus the first, rapid phase of insulin secretion.

**Figure 2.**
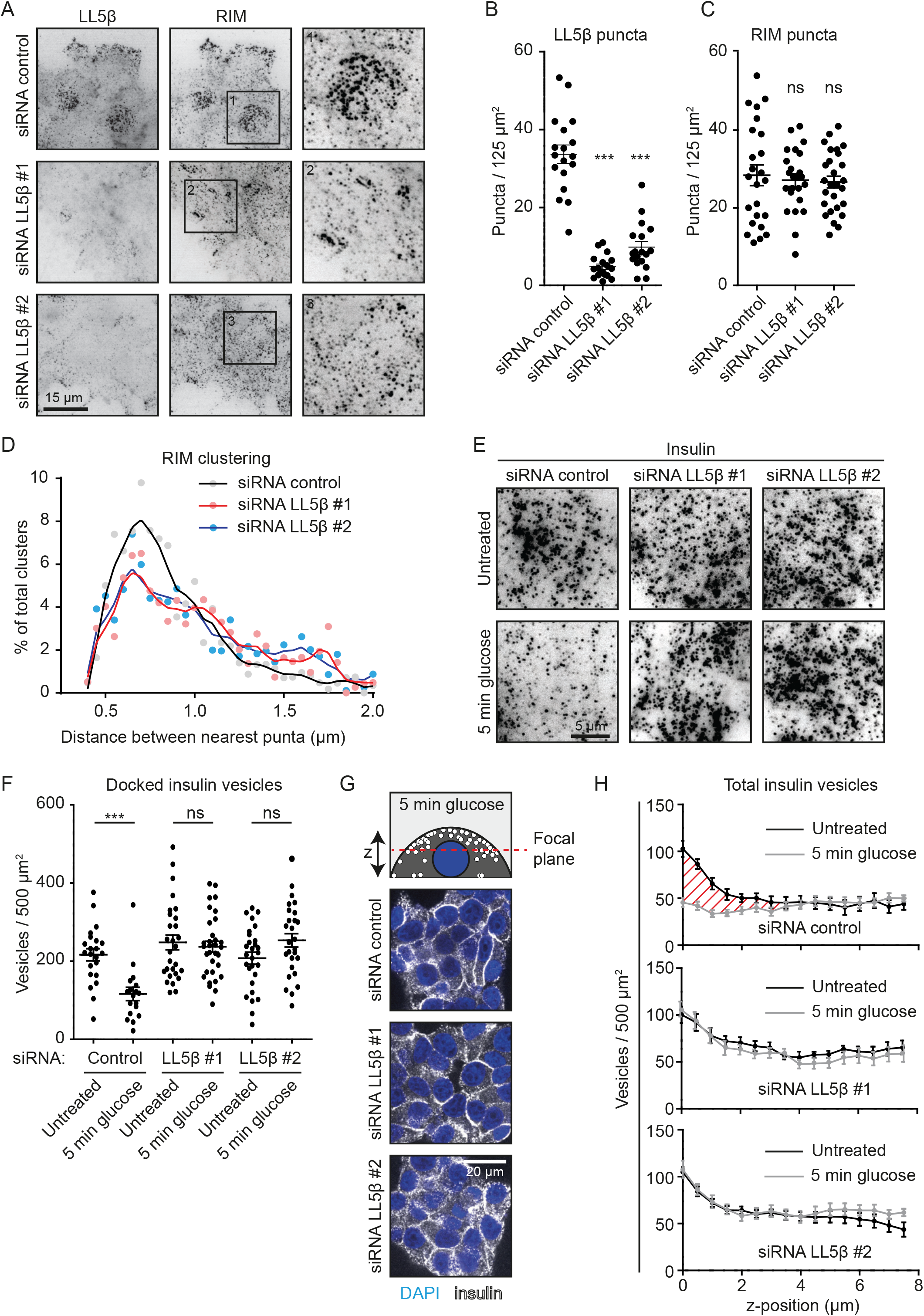
LL5β is required for clustering of insulin docking complexes and insulin release. A. Staining for LL5β and RIM in INS-1E cells transfected with control siRNA or siRNAs against LL5β imaged with TIRFM. B. Quantification of number of LL5β puncta in INS-1E cells treated as in (A). For all conditions, n=18 ROIs (which represent ∼1 cell each); ***p<0.001; one-way ANOVA followed by Dunnett’s post-test. Single data points are plotted. Horizontal line, mean; error bars, S.E.M. C. Quantification of number of RIM puncta in INS-1E cells treated as in (A). Analysis and display as in (B). For all conditions, n=24-27. D. Quantification of RIM clustering in INS-1E cells treated and stained as in (A). Data are plotted as a frequency distribution of distances between nearest puncta. For all conditions, n=24-27 ROIs (which represent ∼1 cell each). Dots represent bin averages; lines represent medium LOWESS (locally weighted scatterplot smoothing) curves. E. Staining for insulin and RIM in INS-1E cells treated as in (A), stimulated with 25 mM glucose as indicated and imaged with TIRFM. F. Quantification of docked insulin vesicles in INS-1E cells treated and stained as in (E). Control, n=18-22 ROIs (which represent ∼4 cells each); LL5β #1, n=28-32; LL5β #2, n=28; ns, no significant difference; ***p<0.001; Mann-Whitney U-test. Single data points are plotted. Horizontal line, mean; error bars, S.E.M. G. Staining for insulin (white) and DNA (blue) in INS-1E cells treated as in (E) and imaged with confocal microscopy. Image focal plane is indicated by red striped line in scheme. H. Quantification of total insulin vesicle distribution along the z-axis in INS-1E cells treated and stained as in (G). For all conditions, n=16 ROIs (which represent ∼4 cells each). Red shaded area indicates secreted insulin fraction at basal side of the cells. Error bars, S.E.M.

### Myosin II activity is required for clustering of insulin secretion complexes around focal adhesions and insulin release

In HeLa cells and keratinocytes, LL5β, liprins, ELKS and KANK1 have been shown to localize around focal adhesions, thereby regulating the CLASP-dependent connection of microtubule plus-ends to the cortex (Bouchet et al., 2016a, Grigoriev et al., 2007, Lansbergen et al., 2006, Stehbens et al., 2014, van der Vaart et al., 2013). This spatial arrangement allows for the coupling between microtubule based transport and localized secretion around focal adhesions (Fourriere et al., 2019, Lansbergen et al., 2006, Stehbens et al., 2014, van der Vaart et al., 2013). Immunofluorescence co-staining of α-tubulin and pFAK visualized spatial proximity of focal adhesions and microtubules in INS-1E cells (Fig. S3A). In addition, co-staining of LL5β and focal adhesion markers in INS-1E and EndoC-βH1 cells showed that LL5β and other cortical proteins localized to the areas adjacent to focal adhesions (Fig. 3A, S3B-E). These cortical complexes, as well as cortical accumulations of insulin granules, were excluded from the regions with dense actin fibers and were located between actin filaments visualized by Single-Molecule Localization Microscopy (SMLM) (Fig. 3B,C).

**Figure 3.**
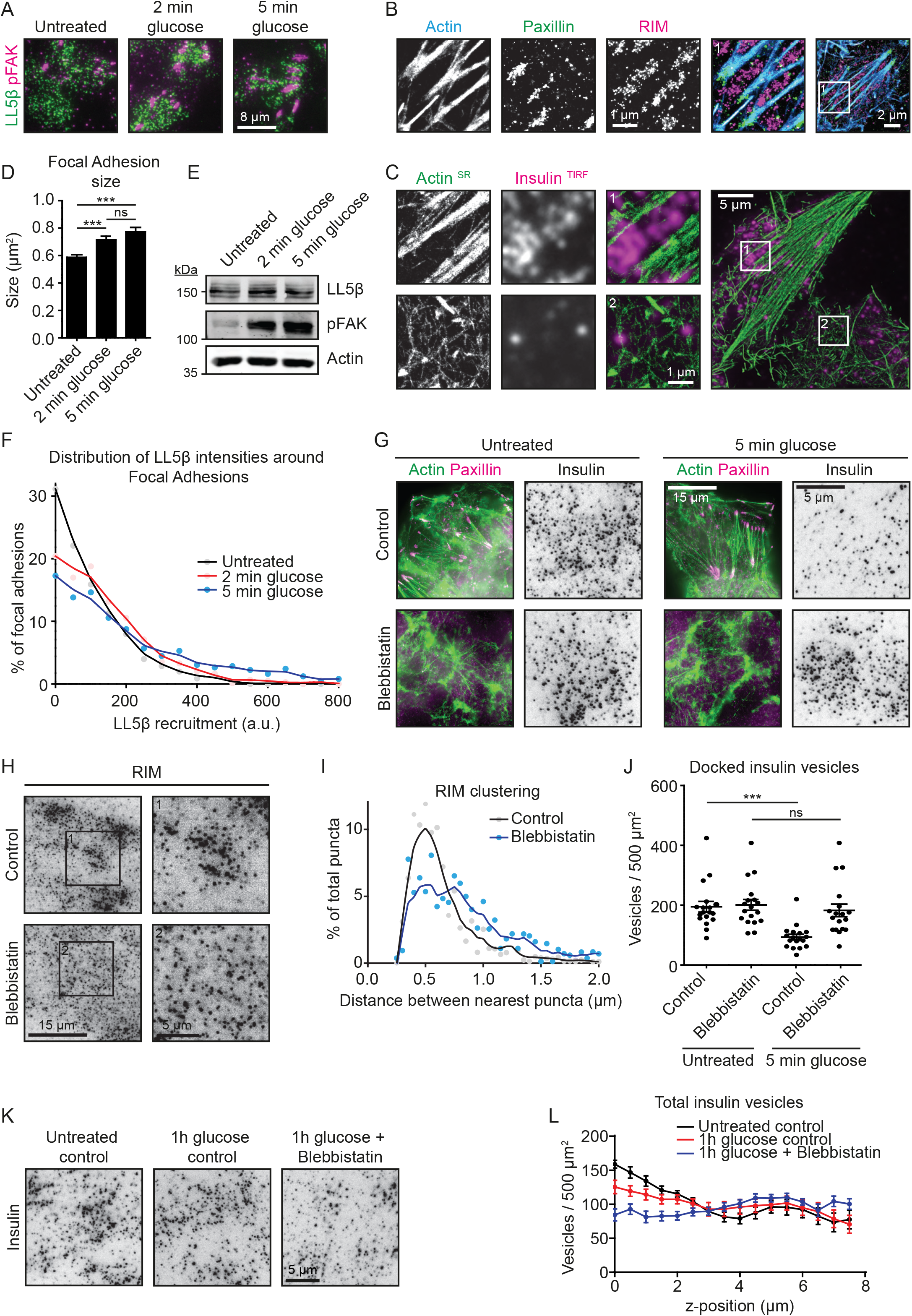
Actomyosin contractility controls the distribution of the cortical insulin docking complexes and insulin release. A. Staining for LL5β (green) and phosphorylated FAK (pFAK, magenta) in INS-1E cells stimulated with 25 mM glucose as indicated and imaged with TIRFM. B. Single-Molecule Localization Microscopy (SMLM) image of actin (blue), paxillin (green) and RIM (magenta) in untreated INS-1E cells. Actin was detected using Peptide-PAINT (Points Accumulation for Imaging in Nanoscale Topography) with LifeAct-mNeonGreen, while paxillin and RIM were detected using DNA-PAINT with antibodies against the endogenous proteins. C. SMLM image of actin (green) detected as in (B) and TIRFM image of insulin (magenta) in INS-1E cells stimulated with 25 mM glucose for 5 min. D. Quantification of focal adhesion size in INS-1E cells treated as in (A). Untreated, n=1040 focal adhesions; 2 min glucose, n=948; 5 min glucose, n=939; ns, no significant difference; ***p<0.001; **p<0.01, one-way ANOVA followed by Tukeýs post-test; error bars, S.E.M. E. Western blot analysis of LL5β and pFAK in INS-1E cells treated as in (A). F. Quantification of LL5β localization relative to focal adhesions in INS-1E cells treated and stained as in (A). Data are plotted as a frequency distribution of LL5β fluorescent intensity (LL5β recruitment) in a 1 µm-broad area around focal adhesions. Only data points exceeding 1.5x LL5β fluorescent background signal where included in plots. Untreated, n=454 focal adhesions; 2 min glucose, n=731; 5 min glucose, n=492. Dots represent bin averages; lines represent medium LOWESS curves. G. Staining for actin (green) and paxillin (magenta), and insulin (grey) imaged with TIRFM in INS-1E cells treated with 50 μM blebbistatin for 1 hour and subsequently stimulated with glucose as indicated. H. Staining for RIM in INS-1E cells treated with 50 μM blebbistatin for 1 hour and imaged with TIRFM. I. Quantification of RIM clustering in INS-1E cells treated and stained as in (H). Analysis and display as in Fig. 2D. For all conditions, n=24. J. Quantification of docked insulin vesicles in INS-1E cells treated and stained as in (G). Analysis and display as in Fig. 2F. For all conditions, n=18. K. Staining for insulin in INS-1E cells treated with 50 μM blebbistatin for 1 hour and subsequently stimulated with glucose as indicated and imaged with TIRFM. L. Quantification of total insulin vesicle distribution along the z-axis in INS-1E cells treated and stained as in (K). Analysis and display as in Fig.2H. For all conditions, n=16.

Work in HeLa cells and fibroblasts previously showed that when focal adhesion formation is suppressed by serum starvation or by inhibition of actomyosin contractility, the clustering of LL5β, liprins, ELKS and KANK1 at the leading cell edges and around focal adhesions is strongly reduced (Bouchet et al., 2016a, Lansbergen et al., 2006). It was also established that glucose stimulation of β-cells leads to activation of focal adhesion kinase (FAK) and remodeling of actin and focal adhesions (Arous & Halban, 2015, Rondas et al., 2011, Rondas et al., 2012). We confirmed these observations: already 2 minutes after glucose stimulation, a significant enlargement of focal adhesions and an increase in the abundance of the phosphorylated, active form of FAK (pFAK) could be detected (Fig. 3A,D,E). Interestingly, glucose stimulation also caused an increase in the abundance of LL5β in the direct vicinity of focal adhesions (Fig. 3A,F). In contrast, when we induced loss of stress fibers and focal adhesion disassembly by inhibiting myosin II with blebbistatin (Fig. 3G), clustering of the components of exocytotic machinery was reduced (Fig. 3H,I), whereas the number of individual puncta was not affected (Fig. S3F). Blebbistatin treatment also inhibited glucose-induced loss of insulin granules from the cell cortex during the first, rapid phase of insulin secretion (Fig. 3G,J). Interestingly, blebbistatin-treated INS-1E cells stimulated for 1 hour with glucose displayed increased loss of insulin vesicles from the cortical area compared to control cells (Fig. 3K,L). This could be explained by the observation that the actomyosin cytoskeleton has an inhibitory effect on the sustained, second-phase insulin secretion (Hammar et al., 2009, Kong et al., 2014). We thus conclude that the role of myosin II in insulin secretion is complex: in the first phase (2-5 min), myosin II is required for the activation of focal adhesion signaling and clustering of the secretion machinery. However, on a longer (1 hour) time scale, loss of cortical actin structures promotes sustained insulin secretion, possibly due to the elimination of the actin cytoskeleton as a cortical barrier for exocytosis, as proposed previously (Arous & Halban, 2015, Arous & Wehrle-Haller, 2017, Kalwat & Thurmond, 2013).

### Visualization of endogenously labelled insulin secretion complexes in mouse pancreatic islets

To confirm that the results obtained in INS-1E cells are valid for endogenous protein complexes, we turned to isolated mouse pancreatic islets. We used islets from wild type mice as well as mice bearing a GFP knock-in in the gene encoding ELKS. In these mice, the GFP coding region with an adjacent neomycin resistance cassette surrounded by two LoxP sites (Lox-Neo-Lox) was inserted directly in front of the first ATG codon in the first coding exon (exon 3) of the *Elks1* gene (Fig. S4A, B). This insertion disrupted the *Elks1* gene, leading to embryonic lethality in homozygous animals, consistent with the previous descriptions of *Elks1* knockout mice (Liu et al., 2014, Wu et al., 2010). When the Lox-Neo-Lox cassette was removed through Cre-mediated recombination, by crossing these mice to a mouse line in which Cre gene is under the control of the cytomegalovirus immediate early enhancer-chicken β-actin hybrid promoter and is expressed in oocytes (Sakai et al., 1997) (Fig. S4B), the resulting *GFP-Elks1* knock-in mice were viable, fertile and displayed no overt defects. Using Western blotting, we observed an upward shift of the ELKS-positive bands in different tissues by ∼30 kDa, as can be expected for GFP fusions (Fig. S4C). These data indicate that the N-terminal fusion to GFP does not perturb the function of the ELKS protein in the mouse, and that the localization of the GFP marker is likely to reflect the endogenous ELKS distribution.

To study ELKS localization and dynamics in mouse β-cells, we isolated pancreatic islets and cultured them on Matrigel-coated coverslips (Fig. S4D). In line with previous publications (Low et al., 2014, Ohara-Imaizumi et al., 2019b, Ohara-Imaizumi et al., 2005), we observed that the endogenously tagged GFP-ELKS preferentially localized along blood vessels within the islets (Fig. 4A,B, Movie S1). In the parts of the isolated islets that adhered to the coverslips, the accumulation of insulin granules and GFP-ELKS were observed in overlapping cortical areas (Fig. 4C). The same was true for the LL5β and RIM detected in islets by immunofluorescence staining (Fig. S4E,F), thus showing that the cortical enrichment patterns of these proteins in mouse pancreatic islets were very similar to those observed in INS-1E cells, human EndoC-βH1 and dispersed human pancreatic islets. Similar localization patterns were also found in human pancreatic tissue, where LL5β and RIM showed colocalization with C-peptide (by-product of insulin production) (Fig. S4G,H).

**Figure 4.**
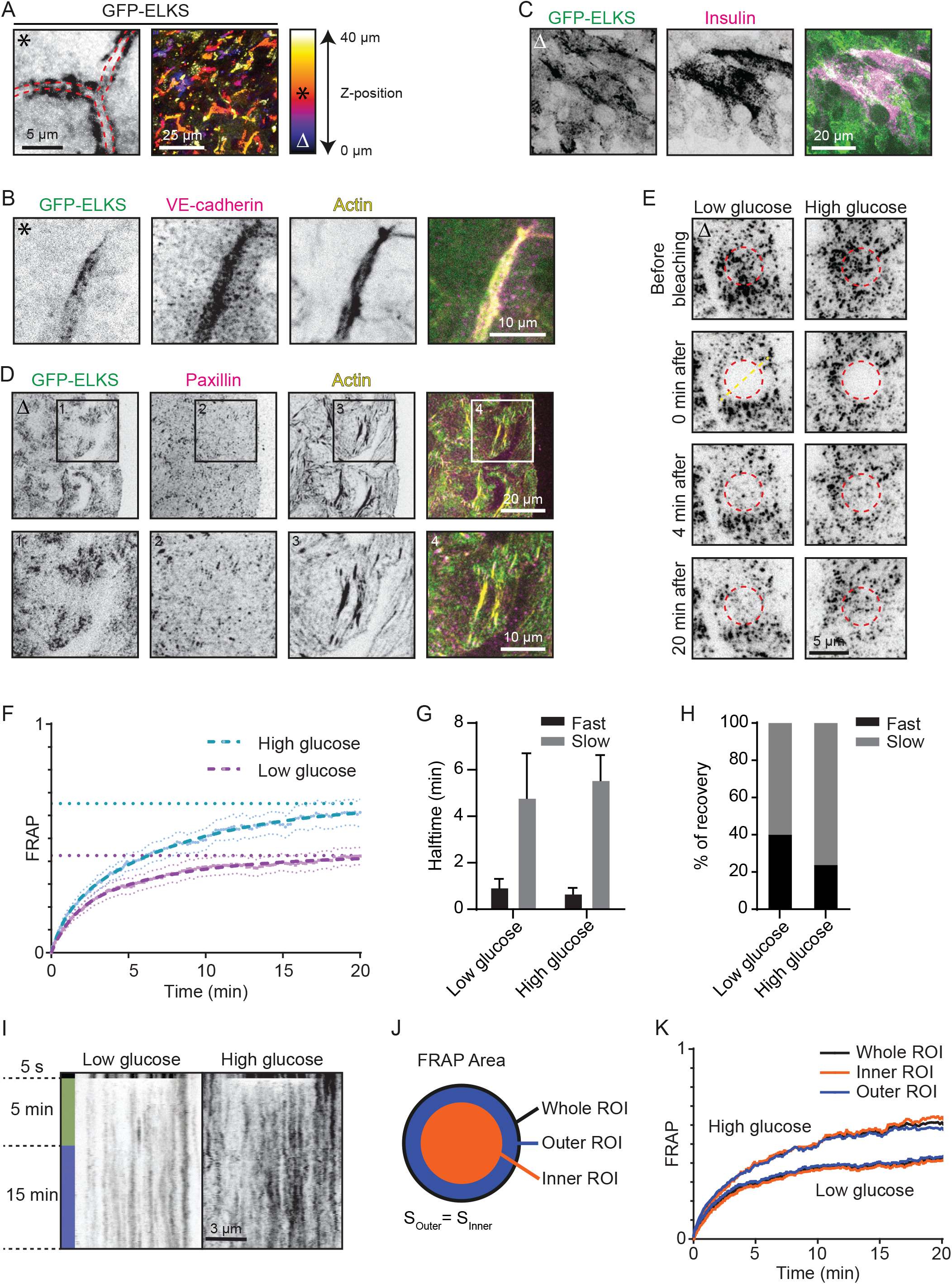
Analysis of the distribution and dynamics of the endogenous cortical insulin secretion complexes in mouse pancreatic islets. A. Localization of GFP-ELKS inside a mouse pancreatic islet imaged with confocal microscopy. Left panel: Red striped lines indicate extracellular space (presumable blood vessel). Right panel: Maximum projection of a z-stack. Image colors indicate z-position (see gradient). Asterix in top left corner of the shown image corresponds to its position in the z-stack (indicated in gradient). B. Staining for VE-cadherin (magenta) and actin (yellow) inside a GFP-ELKS (green)-expressing mouse pancreatic islet imaged with confocal microscopy. Z-position of shown images as in (A). C. Staining for insulin (magenta) in the adherent region of a GFP-ELKS (green)-expressing mouse pancreatic islet imaged with TIRFM. Delta in top left corner of the shown images corresponds to their position in the z-stack (indicated in gradient (A)). D. Staining for paxillin (magenta) and actin (yellow) in the adherent region of a GFP-ELKS (green)-expressing mouse pancreatic islet imaged with TIRFM. Z-position of shown images as in (C). E. FRAP analysis of GFP-ELKS in the adherent region of mouse pancreatic islets stimulated with glucose as indicated and imaged with TIRFM. High glucose (25 mM) was administered 4 hours after low glucose (2 mM) starvation 1 hour prior to photobleaching. Red circles indicate photobleached regions. Z-position of shown images as in (C). F. Average normalized fluorescence intensity recovery and fitted curves (dashed lines) after photobleaching of GFP-ELKS in mouse pancreatic islets treated as in (E) (low glucose, n=15 FRAP areas; high glucose, n=16; N=2 independent experiments). Error lines represent S.E.M, straight horizontal lines represent recovery plateau derived from the fitting. G,H. Fluorescence recovery halftimes (G) and relative fraction (H) of fast and slow exponential components from the fit shown in (F). Error bars represent fitting uncertainty. I. Representative kymographs of GFP-ELKS FRAP along the straight line crossing the center of the bleached area (illustrated as dashed yellow line in (E)). J. Illustration of the original bleaching ROI division into outer and inner areas of equal area for FRAP curves comparison shown in (K). K. FRAP curves corresponding to the full bleaching ROI and its outer and inner areas, as shown in (J).

GFP-ELKS in cultured mouse pancreatic islets displayed a punctate distribution with a characteristic size of an individual spot or a GFP-ELKS cluster being close to the diffraction limit (Fig. 4D). These small clusters were distributed non-homogeneously, often showing local enrichment areas. Such areas often localized around focal adhesions at the base of stress fibers (Fig. 4D), supporting the findings described above.

### Dynamics of GFP-ELKS in pancreatic islets is consistent with binding and unbinding to a scaffold with low mobility

GFP tagging of endogenous ELKS allowed us to measure its dynamics in order to investigate whether the ELKS-containing cortical complexes behave like liquid droplets and whether their turnover is affected by glucose stimulation. To characterize the dynamics of cortical GFP-ELKS puncta at the ensemble level, we performed fluorescence recovery after photobleaching (FRAP) experiments using TIRFM (Fig. 4E, F). For photobleaching, we chose a round area of 5 µm diameter, encompassing multiple clusters (Fig. 4E). Comparison of the FRAP curves showed that both in low glucose and after 1 hour of glucose stimulation, the fluorescence recovery profile could not be satisfactorily fitted by a one-phase association curve, while a much better fitting was obtained using a two-phase curve (Fig. 4F-H). We found that the recovered (exchangeable) fraction of fluorescence in low-glucose conditions was equal to 42%, while a higher recovery of 65% was observed after glucose stimulation (Fig. 4F). Among the parameters describing recovery curves, this shift in the exchangeable/non-exchangeable fractions was the largest, while changes in halftimes of the fast and slow recovering fractions (Fig. 4G) and their relative contributions were relatively minor (Fig. 4H, Supplementary Table 1).

The fluorescence recovery of membrane-associated GFP-ELKS clusters can happen due to their lateral mobility or lateral flux of molecules between them, or through the exchange with the pool of cytoplasmic GFP-ELKS molecules (McSwiggen et al., 2019). If lateral mobility is dominating, faster recovery of the peripheral area of the bleached spot should be observed, due to the flow of unbleached molecules from the periphery. If the proteins predominantly exchange with the cytoplasmic pool, recovery should occur homogeneously throughout the whole bleached area. Kymographs built along the line crossing the center of the bleached round area did not show any noticeable influx of unbleached signal from the periphery (Fig. 4I). To quantify this effect precisely, we divided the bleached spots in two parts of equal areas: peripheral (outer) ring and central (inner) circle (Fig. 4J). The normalized averaged FRAP curves for both areas showed exactly the same behavior as the whole bleached spot (Fig. 4K), suggesting that the recovery happens homogeneously and mostly due to the exchange between cortex-bound and cytoplasmic molecules.

To further confirm the prevalence of this mechanism, we imaged and tracked the mobility of individual GFP-ELKS clusters over the course of 15 minutes (Fig. 5A, Movie S2). The comparison of average mean square displacement (MSD) showed higher cluster mobility upon glucose stimulation (Fig. 5B). In both cases MSD values demonstrated linear dependence on time delay, suggesting diffusive behavior. Linear fits to the MSD plots produced diffusion coefficients of 5.6 x 10^-5^ µm^2^/s for low and 7.5 x 10^-5^ µm^2^/s for high glucose conditions. Although MSD followed a linear dependence, the values of the calculated diffusion coefficients were extremely low and could not account for the observed fluorescence recovery. For example, even in the case of high glucose conditions, the expected FRAP halftime due to the diffusion would be 4 hours (estimated as bleached spot’s area divided by diffusion coefficient (Axelrod et al., 1976)), which is an order of magnitude higher than the observed values (Fig. 4G). Therefore, for the timescale of less than an hour, the contribution of cluster mobility to ELKS dynamics can be excluded from consideration.

**Figure 5.**
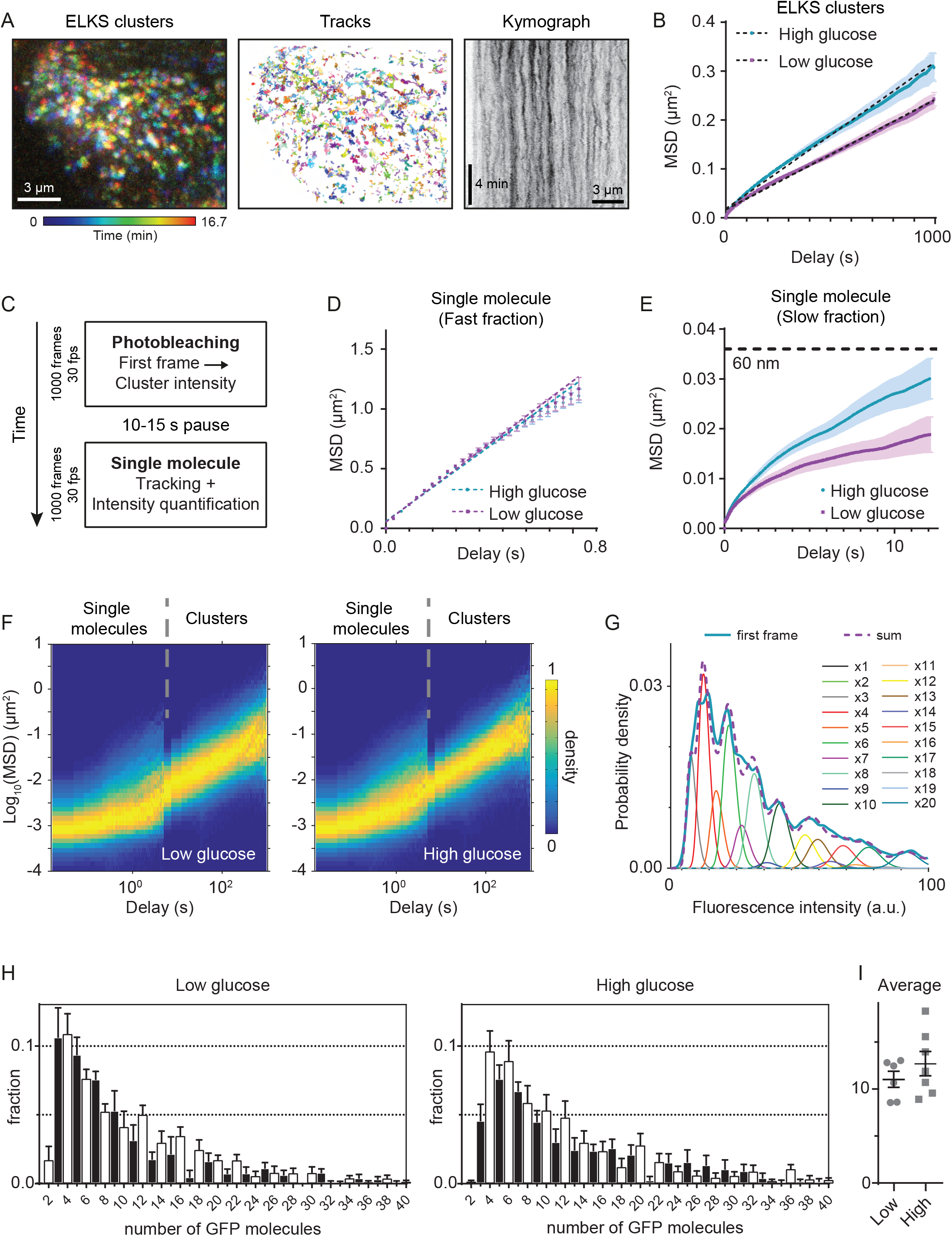
Single molecule analysis of GFP-ELKS cluster motility and stoichiometry in mouse pancreatic islets. A. Representative color coded maximum intensity projection (200 frames, 5 s per frame) of a timelapse movie of GFP-ELKS clusters (left panel) and their corresponding trajectories (middle panel). Right panel shows maximum intensity projection of a stack of kymographs built along horizontal lines of the area shown in the left panel. B. Average MSD of GFP-ELKS clusters trajectories (n=9658 and n=9586 tracks for low and high glucose, N=2 independent experiments). Error bars represent S.E.M. Dashed lines show linear fits MSD(*τ*) = 4D*τ* + σ^2^. Calculated diffusion coefficients are equal to 5.6*10^-5^ µm^2^/s for low and 7.5*10^-5^ µm^2^/s for high glucose conditions. C. Schematic representation of experimental design for observation of single GFP-ELKS molecules. D. Average MSD of fast fraction of single GFP-ELKS molecules trajectories (n=3484 and n=6209 tracks for low and high glucose, N=2 independent experiments). Error bars represent S.E.M. Dashed lines show linear fits MSD(*τ*)=4D*τ* + σ^2^. Calculated diffusion coefficients are equal to 0.37 µm^2^/s for low and 0.35 µm^2^/s for high glucose conditions. E. Average MSD of slow fraction of single GFP-ELKS molecules trajectories (n=4449 and n=7689 tracks for low and high glucose, N=2 independent experiments). Error bars represent S.E.M. Dashed line marks squared displacement of 60 nm, i.e. half the length of GFP-ELKS molecule (119 nm (Sala et al., 2019)). F. Combined heatmap (3D histogram) of MSD values for the slow fraction of single GFP-ELKS molecules and clusters (separated by gray dashed lines). Same datasets as in (E) and (B). Histogram values are normalized by the maximum value of each column, corresponding to each time delay bin. G. Representative probability density of pre-bleach GFP-ELKS clusters intensities (measured at the first frame, thick blue line, one FOV, 1095 clusters) fitted to a weighted sum of *N*-mers of GFP (thick dashed magenta line). The weighted probability densities of individual GFP *N*-mers intensities are plotted as thin lines. H. Quantification of GFP-ELKS cluster stoichiometry. Averaged histograms of weights of *N*-mers of GFP determined from the fitting to the GFP-ELKS clusters intensities (n=6 FOVs for low and n=7 FOVs for high glucose, N=2 independent experiments). Error bars represent S.E.M. Weighted mean values of GFP molecules per cluster for each FOV are shown in the inset (11.0 ± 2.1 for low and 12.7 ± 3.5 for high glucose condition, average ± S.E.M.). I. Average number of GFP-ELKS molecules present in a cluster.

The formation of (relatively) immobile clusters from the molecules present in a cytosolic pool can happen through their binding/unbinding to a scaffold formed by other proteins. Alternatively, as suggested by recent publications (Liang et al., 2021, Sala et al., 2019), ELKS cluster formation could occur spontaneously due to liquid-liquid phase separation. To discern among these two mechanisms, we measured the behavior of GFP-ELKS at the level of individual molecules. We performed a set of recordings on the previously unexposed area of islets with high laser power and high frame rate (Fig. 5C). During the first acquisition, we photobleached all fluorescent signal in the field of view, including bright immobile GFP-ELKS clusters at the cortex. After a short (10-15 s) recovery period, we performed a second high-frame-rate acquisition to record the behavior of single GFP-ELKS molecules (Fig. 5C). Improved contrast after the bleaching of bright immobile spots allowed us to observe the presence of two populations of single molecules: a slow, relatively immobile pool and particles that were rapidly diffusing in and out the field of view (Movie S3). We observed multiple events of transitions between these two populations (Movie S4), confirming the exchange between clusters and cytosolic pool. Substantial photobleaching (caused by high laser power required for the observation of fast-moving molecules) precluded precise quantification of corresponding rate constants of transitions between these fractions. Therefore, we focused on the characterization of motility of GFP-ELKS molecules. Our first attempt of tracking single molecules led to multiple mistakes at the step of linking detections to trajectories (Fig. S5A, Movie S3, left panels). Automatic linking methods based on the nearest neighbor search generated trajectories that were composed from interspersed fragments of tracks from both populations (Movie S3, left panels). This problem hindered the analysis of trajectories. As can be seen from the MSD density plot (Figure S5D, left panel), two separate particle populations are present, manifesting themselves as two separate dense areas/shapes, where the higher one corresponds to the fast-and the lower one to the slow-moving population. The average duration of tracks, measured as the length of each area along the *x* axis, appeared longer for the fast-moving population. This was in contradiction with the visual inspection of acquired movies, where fast-moving particles rapidly enter and leave the imaging volume, while the slow molecules remain in the field of view for much longer periods. To overcome this analysis problem, we applied a temporal median filter with the sliding window of 15 frames (∼0.5 s) to each pixel of the original movie (Hoogendoorn et al., 2014, Masucci et al., 2021). As a result, we obtained two movies with clearly separated timescales, where each of the two populations can be tracked independently (Fig. S5B,C, Movie S3). In this case, MSD density plot correctly reflected shorter duration of fast-moving particle trajectories in comparison to their slow-moving counterparts and demonstrated a two order difference in the motility between them (Figure S5D, right).

The average MSD curves of the fast fraction of single GFP-ELKS molecules did not depend on the glucose exposure (Fig. 5D). Linear fits provided similar diffusion coefficient values of 0.37 µm^2^/s for low and 0.35 µm^2^/s for high glucose conditions. This allowed us to attribute the fast-moving population to the GFP-ELKS molecules freely diffusing in the cytosol. In contrast, the mobility of the slow fraction was dependent on glucose exposure (Fig. 5E) in a similar way to the mobility of clusters (Fig.5B). More importantly, in this case the average displacement of GFP-ELKS molecules on the timescales of up to 10 seconds did not exceeded 60 nm (Fig. 5E). Given that the size of ELKS dimer determined by EM data is 119 nm (Sala et al., 2019), this means that the individual molecules are essentially immobile within the cluster. To combine the MSD data of GFP-ELKS single molecules and cluster trajectories on different timescales, we generated “stitched” MSD probability heatmap (Fig. 5F). We observed that the mode (peak) of MSD distribution of single molecules continuously transitioned to the mode of clusters for each glucose concentration (Fig. 5F). Therefore, we conclude that the slow movement of GFP-ELKS single molecules can be explained by movement of clusters as a whole, rather than the diffusion of individual molecules within them. Lack of mobility of individual molecules combined with the slow diffusion of the clusters speaks in favor of the hypothesis that the ELKS molecules are bound to a low-mobility scaffold, being in binding/unbinding equilibrium with cytosolic ELKS molecules. If each cluster would represent a separate liquid phase/droplet, one would expect to observe diffusion of individual molecules within it.

To characterize the size of individual GFP-ELKS clusters, we applied a quantitative fluorescence approach (Fig. 5G,H). Using single molecule intensity in the last frame of the tracks, before complete photobleaching, we estimated fluorescence intensity distribution of single GFP fluorophores. Using this distribution, we constructed GFP N-mer distributions corresponding to intensities of oligomers with increasing number N of GFP molecules (Moertelmaier et al., 2005). Initial, unbleached intensity of GFP-ELKS clusters (at the first frame) was fitted to a sum of these distributions with different weights, which served as fitting parameters (Fig. 5G). We found that oligomers with an even number of GFP molecules were prevalent in the intensity distributions (Fig. 5H), in accordance with a previous report showing that ELKS forms coiled-coil homodimers (Sala et al., 2019). The weighted average number of GFP molecules per cluster did not change significantly and was measured as 11.0 ± 2.1 for low glucose and 12.7 ± 3.5 for high glucose condition (Fig. 5I). In both cases the most abundant oligomer fraction contained four GFP molecules (Fig. 5H), corresponding to just two ELKS homodimers. It is highly improbable that a stable molecular assembly with such a low number of molecules can be formed as a result of liquid-liquid phase separation. Therefore, our stoichiometry data analysis strongly favors the idea that GFP-ELKS clusters form by binding to relatively immobile scaffolds.

## Discussion

In this study we showed that the sites of insulin secretion in pancreatic β-cells combine molecular components of the neuronal CAZ, controlling the very rapid Ca^2+^-regulated neurotransmitter secretion, and the cortical platforms responsible for constitutive exocytosis at the leading edges and around focal adhesions in migrating non-neuronal cells (Fig. 6). Presence of non-neuronal components such as LL5β and KANK1 explains the clustering of exocytotic sites in cortical regions where β-cells form integrin-based adhesions to the extracellular matrix - the basement membrane surrounding blood vessels.

**Figure 6.**
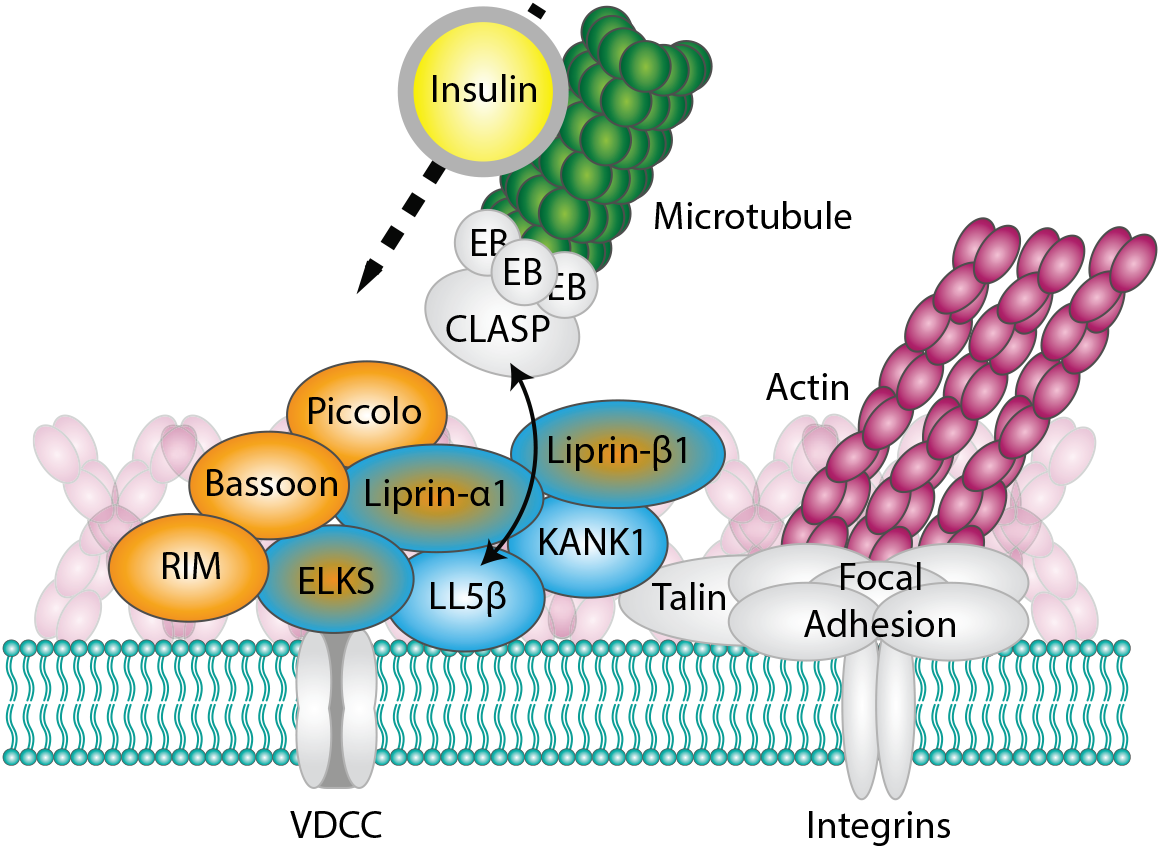
Schematic representation of secretory sites in pancreatic β-cells. CAZ-specific components are indicated in orange, components that are not present in neurons, in blue, and components shared between neuronal and non-neuronal cells, by both colors.

Previous work has shown that the depletion of LL5β perturbs clustering of its cortical partners (Lansbergen et al., 2006, van der Vaart et al., 2013), and we found that the same is true in β-cells. Depletion of LL5β affected the first, rapid phase of insulin release. Enrichment of LL5β and the microtubule-stabilizing protein CLASP1 at exocytotic sites suggests that these proteins contribute to efficient microtubule connection to the cortex, which could help to balance microtubule-based delivery and withdrawal of insulin granules from secretory platforms, as proposed previously (Bracey et al., 2020, Heaslip et al., 2014, Ho et al., 2020, Tabei et al., 2013, Varadi et al., 2003, Yuan et al., 2015, Zhu et al., 2015). In this way, secretory sites in β-cells are reminiscent of LL5β- and CLASP-containing podosome-like structures (“synaptic podosomes”), which provide a route for microtubule-based delivery and accumulation of acetylcholine receptors at neuromuscular junctions in skeletal muscle cells (Basu et al., 2015, Kishi et al., 2005, Proszynski et al., 2009, Proszynski et al., 2013, Schmidt et al., 2012). However, although microtubules are in tight contact with the secretion complexes and insulin granules (Muller et al., 2021, Yuan et al., 2015), it is unlikely that the loss of LL5β perturbs the rapid phase of insulin exocytosis by reducing microtubule density. First, this secretion phase depends on insulin granules that are already docked at the cortex and do not need to be transported along microtubules. Second, recent evidence suggests that dense microtubules at the cortex attenuate rather than promote insulin secretion by controlling vesicle movement away from the plasma membrane (Bracey et al., 2020, Ho et al., 2020, Zhu et al., 2015). Therefore, we favor the idea that LL5β stimulates insulin secretion by directly enhancing clustering of CAZ components such as RIM, ELKS and liprins, which in turn control, either directly or indirectly, vesicle tethering, connections to the core fusion machinery such as SNAREs and their regulators, and the activation of voltage-gated calcium channels (reviewed in (Gundelfinger & Fejtova, 2012, Ohara-Imaizumi et al., 2019a, Sudhof, 2012)).

Tight clustering of proteins participating in secretion is a hallmark of neuronal CAZ, and recent work in worms showed during synapse development the worm homologues of ELKS and liprin undergo LLPS and then solidify (McDonald et al., 2020). Moreover, different mammalian CAZ components can undergo LLPS in vitro or when overexpressed in cells (Emperador-Melero et al., 2021, Liang et al., 2021, McDonald et al., 2020, Sala et al., 2019, Wu et al., 2019, Wu et al., 2021). This raised the question of whether LLPS plays a role in organizing secretion machinery in β-cells. Our previous work in non-neuronal cells showed that LL5β, KANK1 and liprins are recruited to the plasma membrane independently of each other and display their own characteristic turnover, whereas their multivalent interactions with each other and additional membrane-unbound scaffolds such as ELKS promotes their clustering (Bouchet et al., 2016a, Lansbergen, 2006, van der Vaart et al., 2013). A similar picture emerges in β-cells: all investigated cortical proteins appear as closely apposed but not fully overlapping membrane-associated foci, which, based on the imaging of endogenously tagged ELKS, contain only a few protein copies that exchange with the cytoplasmic pool and undergo very slow lateral diffusion. The half-time of the slower component of recovery of endogenously tagged ELKS in pancreatic islets (∼5 min) was very similar to that of mildly overexpressed ELKS or liprin-α1 in HeLa cells (van der Vaart et al., 2013), suggesting that ELKS behaves similarly in other non-neuronal cells. Formation of liquid droplets of ELKS in non-neuronal cells (Sala et al., 2019) thus appears to be a consequence of its overexpression. It is still possible that CAZ in neurons does form by LLPS, for example, because the expression of key players might be higher in neurons than in pancreatic cells, or because more LLPS-prone protein isoforms (e.g. liprin-α3 rather than liprin-α1 (Emperador-Melero et al., 2021)) are expressed. Single molecule imaging of endogenously tagged mammalian CAZ components in neurons will be needed to address this possibility.

An important distinguishing feature of the secretion sites in β-cells compared to neuronal CAZ is their association with focal adhesions. Integrin activation provides a spatial cue for targeting insulin secretion to blood vessels ((Gan et al., 2018); reviewed in (Arous & Halban, 2015, Arous & Wehrle-Haller, 2017)). Previous work showed that elevated glucose levels lead to enlargement of focal adhesions (Arous & Halban, 2015, Arous & Wehrle-Haller, 2017), and here we found that this is accompanied by enhanced clustering of secretion complexes around adhesions and their increased lateral mobility and turnover. The biochemical basis of this regulation needs to be elucidated and may involve alterations in the state of focal adhesion components, such as conformational changes in talin promoting KANK1 recruitment (Yu et al., 2019), or activation of FAK (Rondas et al., 2011, Rondas et al., 2012) and other integrin-linked signaling pathways.

Formation and regulation of focal adhesions critically depends on actomyosin contractility. Here, we showed that inhibition of myosin II, which causes focal adhesion disassembly, had a strong negative effect on the clustering of secretory complexes and the first phase of insulin release from β-cells. This effect represents one of the multiple and complex roles that actin and myosin II play in insulin secretion. Actin filaments in β-cells are seen as tracks for vesicle transport and as a barrier for secretion (Arous & Halban, 2015, Arous & Wehrle-Haller, 2017, Kalwat & Thurmond, 2013). We observed that the exocytotic sites and cortically docked insulin granules form a complementary pattern with the actin filaments, consistent with the idea that a dense cortical actin meshwork might inhibit secretion. On the other hand, myosin II is universally recognized as an important positive regulator of secretion, which acts at different levels, from indirect reorganization of the cortical cytoskeleton to direct contribution to the release of vesicle content during secretion (Arous & Halban, 2015, Gutierrez et al., 2018, Rousso et al., 2016).

Finally, it is important to note that the interactions between actomyosin, integrin adhesions and secretory machinery are subject to intricate feedback mechanisms. For example, in migrating cells, LL5β and liprins can regulate integrin activation and focal adhesion turnover (Asperti et al., 2009, Asperti et al., 2010, Stehbens et al., 2014). Since integrin-based adhesion to extracellular matrix plays a crucial role in β-cell survival (Bosco et al., 2000, Hammar et al., 2004, Weber et al., 2008), understanding the interplay between secretory machinery, integrin adhesion and the cytoskeleton is important for developing new strategies to treat type 2 diabetes.

## Materials and Methods

### Cell lines

INS-1E cells (Merglen et al., 2004) (kind gift of B. Guigas, LUMC) were cultured in Roswell Park Memorial Institute (RPMI) medium 1640 (Gibco, Life Technologies) supplemented with 10% fetal calf serum (FCS) (GE Healthcare Life Sciences), 10 mM HEPES pH 7.4, 1 mM Sodium Pyruvate (Gibco, Life Technologies), 55 μM β-mercaptoethanol (Gibco, Life Technologies) and 11 mM glucose. The human insulinoma cell line EndoC-βH1 cells (Ravassard et al., 2011) obtained from Univercell-Biosolutions, was cultured in complete medium (low glucose DMEM (Gibco, 1g/L glucose), 2% Albumin (Sanquin Bloodbank, the Netherlands), 10 mM nicotinamide (prepared by the LUMC pharmacy), 5.5 g/mL Human Transferrin (Sigma-Aldrich), 0.5 mg/mL Selenite (Sigma-Aldrich), 10 mL Penicillin (100U/mL) /Streptomycin (100g/mL), with fresh β-mercaptoethanol (Sigma-Aldrich) added when culturing (to a final concentration of 0.05 mM)) in a humidified atmosphere with 5% CO_2._ The cells were routinely checked for mycoplasma contamination using the MycoAlert^TM^ Mycoplasma Detection Kit (Lonza).

### Mouse pancreatic islets

Both male and female adult 3-6-month-old mice were sacrificed by cervical dislocation for the isolation of tissues and primary cultures. All animal experiments were performed in compliance with the institutional guidelines for the welfare of experimental animals approved by the Animal Ethical Review Committee (DEC 2014.I.03.020) of Utrecht University, the Netherlands.

The isolated pancreas was rinsed in PBS, cut into small pieces with surgical blades and incubated in 3 mg/ml collagenase (C9263, Sigma) dissolved in glucose-free RPMI 1640 (Gibco, Life Technologies) in a total volume of 3 ml/pancreas for 20 min at 37°C. During incubation, solution was shaken rigorously. Next, 6 ml of cold RPMI 1640 supplemented with 10% FCS, 10 mM Hepes pH 7.4, 1 mM Sodium Pyruvate, 55 μM β-mercaptoethanol and 11 mM glucose was added to the solution, followed by two wash steps using centrifugation (5 min, 200 x g, 4°C). After washing, the isolated islets were resuspended in 5 ml RPMI 1640 supplemented as described above, handpicked using a pipette and transferred to new culture dishes until almost no exocrine tissue was left. Finally, the islets were transferred to culture dishes containing Matrigel-coated coverslips. Coverslips where coated with 388 μg/ml Matrigel (Corning) in phosphate-buffered saline (PBS) for 2 hours at 37°C. Isolated islets were cultured in RPMI 1640 supplemented as described for the rat INS-1E cell line until islets were fully attached to coated coverslips.

### Human pancreatic islets

Human islet isolations from cadaveric human organ donors were performed in the Good Manufacturing Practice facility of the Leiden University Medical Center according to the method used in the center for the procurement of clinical-grade material (Nijhoff et al., 2016). Islets were used for research only when they could not be used for clinical purposes; research consent had been obtained according to national laws.

Islets were cultured in CMRL 1066 medium (Corning, 5.5 mmol/L glucose) containing 10% FCS (Bodinco), 20 mg/mL ciprofloxacin (Fresenius), 50 mg/mL gentamycin (Lonza), 2 mM l-glutamine (Lonza), 10 mM HEPES (pH 7.21, Lonza Lonza), and 1.2 mg/mL nicotinamide (prepared by the LUMC pharmacy) in a humidified atmosphere with 5% CO_2_. Islets were dispersed into single cells by adding 0.025% trypsin solution containing 10 mg/mL DNase (Pulmozyme, Genentech) at 37°C for 6-8 minutes.

### Immunofluorescence staining of fixed samples

For immunofluorescence staining experiments, cells or pancreatic islets were seeded or attached on coverslips coated with fibronectin (INS-1E, EndoC-βH1), ECM gel (EndoC-βH1, Engelbreth-Holm-Swarm murine sarcoma), Matrigel (mouse pancreatic islets) or poly-L-lysine (dispersed human pancreatic islets). Cells or pancreatic islets were fixed with either 4% paraformaldehyde for 20 min at room temperature (staining for insulin, LL5β, RIM1/2, Bassoon, paxillin, glucagon, VE-cadherin, phalloidin) or −20°C MeOH for 10 min (staining for LL5β, ELKS, liprin-α1, liprin-β1, KANK1, RIM1/2, Bassoon, CLASP1, pFAK) followed by permeabilization with 0.15% Triton X-100 for 2 min. Next, samples were blocked with 1% bovine serum albumin (BSA) diluted in phosphate buffered saline (PBS) supplemented with 0.05% Tween-20 for 45 min at room temperature and sequentially incubated with primary antibodies for 1 hour at room temperature (pancreatic islets: overnight at 4 °C) and fluorescently labeled with secondary antibodies for 45 min at room temperature (pancreatic islets: 1-3 hours). Finally, samples were washed, dried and mounted in DAPI-containing Vectashield (Vector laboratories).

For human tissue staining experiments, pancreatic tissue biopsies (1 cm x 1 cm x 1 cm max) were fixed in 4% Formalin (Klinipath) for 24-48 hours. After fixation, tissue was placed in 70% ethanol, before moving to dehydration in an ascending series of ethanol (10-100%), followed by xylene and paraffin. Paraffin blocks were cut into 4 µm sections (Leica RM2255 Microtome). Sections were rehydrated and antigen retrieval was performed by heating slides in citrate buffer (pH 6.0) using a pressure cooker. Sections were blocked with 5% normal donkey serum (in 0,3% Triton-100 in PBS), stained with primary antibodies (LL5β, RIM) overnight at 4°C followed by 1-hour incubation with fluorescently labeled secondary antibodies. Subsequently, sections were incubated for 1 hour at room temperature with a primary antibody against C-peptide (Abcam), followed by 1-hour incubation with corresponding fluorescently labeled secondary antibody. Finally, samples were incubated for 3 minutes with Hoechst for nuclear staining and slides were mounted with Prolong gold (ThermoFisher).

For immunofluorescence staining prior to SMLM, cells were extracted and fixed in 0.5% Triton-X100 and 4% PFA in cytoskeleton buffer (10 mM MES, 150 mM NaCl, 5 mM MgCl_2_, 5 mM EGTA, 5 mM Glucose, pH 6.1) for 15 minutes at room temperature. Next, cells were blocked in 1% BSA diluted in PBS supplemented with 0.05% Tween-20 for 45 min at room temperature and sequentially stained with primary antibodies overnight at 4°C. After three washes in PBS, cells were incubated with either a fluorescently labeled secondary antibody or a DNA-sequence conjugated secondary antibodies (Ultivue) for 1.5 hours at room temperature. Coverslips were mounted in a Ludin chamber.

### Co-immunoprecipitation from INS-1E cell extracts

For co-immunoprecipitation, proteins were extracted from INS-1E cells in a Triton X-100 lysis buffer (20 mM Tris-Cl, pH 7.5, 150 mM NaCl, 0.5 mM EDTA, 1 mM DTT, 1% Triton X-100, and a protease inhibitor cocktail (Roche)) with gentle rotation for 30 minutes at 4 °C. All further steps were also executed at 4 °C. The sample was centrifuged for 20 min at 16,100 x g to obtain the supernatant, which was further incubated with Affi-prep Protein A beads (Biorad) for preclearing for 30 minutes. The sample was centrifuged for 10 minutes at 16,100 x g and the supernatant was further incubated with rabbit control serum or anti-ELKS antibody (Proteintech) for 1 hour. After adding Affi-prep Protein A beads, the sample was further incubated for 1 hour with rotation. The beads were washed three times before the proteins were eluted by boiling them in an SDS-PAGE sample buffer. The samples were subjected to SDS-PAGE followed by Western Blot for analysis.

### siRNA transfection

Lipofectamine RNAiMAX (Invitrogen) was used to transfect INS-1E cells with 20 nM siRNAs. Corresponding experiments were performed 72 hours after siRNA transfection.

### Glucose and drug treatment of INS-1E

Before the induction glucose stimulated insulin secretion, INS-1E cells were glucose starved by culturing the cells for a minimum of 4 hours in culture medium as described above supplemented with 2 mM glucose (termed untreated or low glucose). Glucose stimulated insulin secretion was induced by culturing glucose starved INS-1E cells in the culture medium as described above supplemented with 25 mM glucose (termed high glucose). Cells were treated with 50 μM blebbistatin for 1 hour prior to fixation or live cell imaging.

### Generation of GFP-ELKS mouse line

The knockout targeting construct was generated by inserting GFP fused in frame with an N-terminal peptide that can be biotinylated by the biotin ligase BirA (de Boer et al., 2003) and a neomycin resistance cassette flanked by *loxP* sites in front of the start codon of the mouse *Elks1* gene into exon 3 using a PCR-based strategy. The construct was linearized and electroporated into IB10 embryonic stem (ES) cells, which were cultured in BRL-cell conditioned medium as described previously (Hoogenraad et al., 2002). Targeted ES cells were further selected with G418 (200 µg/ml; Life Technologies) for neomycin resistance, and individual clones were picked and expanded. Genotyping by PCR was performed to check for the positive clones, which were subsequently injected in blastocysts obtained from C57Bl/6 females. Male chimera mice were mated with C57BL/6 females to transmit the ELKS knockout allele to the germline. The generation of the *Elks1* knockout mice was performed in compliance with the institutional guidelines and approved by the Animal Ethical Review Committee (DEC) of the Erasmus Medical Center, Rotterdam, the Netherlands.

To obtain GFP-ELKS knock-in mice, heterozygous Elks1 knockout mice were crossed with mice in which Cre gene is under the control of the cytomegalovirus immediate early enhancer-chicken β-actin hybrid promoter and is expressed in oocytes (Sakai & Miyazaki, 1997). Mice were genotyped by PCR.

### Preparation of mouse tissue extracts

Mouse tissues were dissected and placed in ice-cold PBS, pH 7.4. Samples were weighted and homogenized in ice-cold homogenization buffer (150 mM NaCl, 50 mM Tris, 0.1% v/v SDS, 0.5% v/v NP-40 pH8, 1x complete protease inhibitor, Roche) with stainless metal beads (Qiagen) using the TissueLyser II (Qiagen) for 30 minutes. Tissue lysates were then centrifuged at 16,000 x g at 4 °C for 1 hour. Supernatant was used for Western blotting.

### Fluorescence microscopy

#### Widefield microscopy

Fixed and stained INS-1E cells were imaged using widefield fluorescence illumination on a Nikon Eclipse 80i upright microscope equipped with a CoolSNAP HQ2 CCD camera (Photometrics), an Intensilight C-HGFI precentered fiber illuminator (Nikon), ET-DAPI, ET-EGFP and ETmCherry filters (Chroma), controlled by Nikon NIS Br software and using a Plan Apo VC 23 100x NA 1.4 oil, Plan Apo VC 60x NA 1.4 oil or a Plan Fluor 20x MI NA 0.75 oil objective (Nikon). For presentation, images were adjusted for brightness using ImageJ 1.50b.

#### TIRF microscopy

Fixed and stained INS-1E cells, EndoC-βH1, dispersed human pancreatic islets and live GFP-ELKS expressing pancreatic islets were imaged using TIRFM performed on an inverted research microscope Nikon Eclipse Ti-E (Nikon) with a perfect focus system (PFS) (Nikon), equipped with a Nikon CFI Apo TIRF 100 × 1.49 N.A. oil objective (Nikon), Evolve 512 EMCCD (Photometrics) or Prime BSI camera (Photometrics) or CoolSNAP HQ2 CCD camera (Roper Scientific) and controlled with the MetaMorph 7.7.11.0 software (Molecular Devices). Images were projected onto the chip of an Evolve 512 camera with an intermediate lens 2.5× (Nikon C mount adapter 2.5×) or onto a CoolSNAP HQ2 or a Prime BSI chip without the lens. In all cases the final magnification was equal to 0.065 μm/pixel. To keep cells at 37°C a stage top incubator INUBG2E-ZILCS (Tokai Hit) was used. For excitation, 491 nm 100 mW Stradus (Vortran), 561 nm 100 mW Jive (Cobolt) and 642 nm 110 mW Stradus (Vortran) lasers were used. The microscope was equipped with an ET-GFP 49002 filter set (Chroma) for imaging of proteins tagged with GFP, an ET-mCherry 49008 (Chroma) and an ET-405/488/561/647 filter set. For presentation, images were adjusted for brightness using ImageJ 1.50b.

#### Spinning disk microscopy

Fixed and stained INS-1E cells, dispersed human pancreatic islets and isolated mouse pancreatic islets were imaged using confocal fluorescence illumination on a Nikon Eclipse Ti microscope equipped with a perfect focus system (PFS, Nikon), a spinning-disc-based confocal scanner unit (CSU-X1-A1, Yokogawa), an Evolve 512 EMCCD camera (Photometrics) attached to a 2.0× intermediate lens (Edmund Optics), a super-high-pressure mercury lamp (C-SHG1, Nikon), a Roper Scientific set of Stradus 405-nm (100 mW, Vortran), Calypso 491-nm (100 mW, Cobolt) and Jive 561-nm (100 mW, Cobolt) lasers, a set of ET-BFP2, ET-EGFP, ET-mCherry and ET-EGFP-mCherry filters (Chroma) for wide-field fluorescence observation, a set of ET460/50m, ET525/50m or ET535/30m (green), ET630/75m (red) and ET-EGFP/mCherry filters (Chroma) for spinning-disc-based confocal imaging and a motorized stage MS-2000-XYZ with Piezo Top Plate (ASI). The microscope setup was controlled by MetaMorph 7.7.11.0 software. Images were acquired using Plan Fluor 10× NA 0.3 air, Plan Fluor 20× MI NA 0.75 oil, Apo LWD λS 40x NA 1.15 water, Plan Apochromat λ 60× NA 1.4 oil and Plan Apo VC 60× NA 1.4 oil objectives. Images are presented as single plane images or maximum projections of 0.5-μm-step z-scans and adjusted for brightness using ImageJ 1.50b.

Fixed and stained human pancreatic tissue sections were examined using a commercially available DragonFly200 spinning disk system (Andor) on a DMi8 microscope (Leica) with Plan Apo 40× NA 1.3 and Plan Apo 63× NA 1.4 oil immersion objectives. Microscope setup was controlled by Fusion software (Andor), and images were taken with a Zyla sCMOS camera (Andor).

#### STED microscopy

Super-resolution imaging of cortical proteins in INS-1E cells was performed using gated STED modality on a Leica TCS SP8 STED 3X SMD confocal microscope, using spectroscopic detection with HyD hybrid detector. For the illumination we used a fully tunable supercontinuum white light laser WLL (470 to 670 nm) and 592 nm, 660 nm and 775 nm STED depletion lasers. Images were acquired using a HC PL APO CS2 100×/1.40 oil objective. Alexa-Fluor-488-conjugated antibodies were excited with the 488 nm WLL (80MHz) and depleted with the CW 592 nm STED depletion laser. Alexa-Fluor-594-conjugated antibodies were excited with the 594 nm super continuum white laser (80MHz) and depleted with the 775 nm STED pulsed depletion laser. For presentation, images were adjusted for brightness using ImageJ 1.50b.

#### Single molecule localization microscopy

Single Molecule Localization Microscopy (SMLM) of fixed INS-1E cells was performed using a Nikon Ti-E microscope with a 100x Apo TIRF oil immersion objective (NA1.49) and a Perfect Focus System 3. Excitation was performed through a custom illumination pathway with a Lighthub-6 (Omicron) laser containing a 638 nm laser (BrixX 500 mW multimode, Omicron), a 488 nm laser (Luxx 200 mW, Omicron), and a 405 nm laser (Luxx 60 mW, Omicron) and optics to tune the incident angle for evanescent wave or oblique illumination. Signal was detected with a sCMOS camera (Hamamatsu Flash 4.0v2). For imaging of actin and insulin, first a widefield image of insulin was obtained. Then, a low concentration of LifeAct-mNeonGreen was added such that single molecules could be observed and ∼25000 frames were acquired at 60 ms exposure to reconstruct a super resolved image (Schatzle et al., 2018, Tas et al., 2018). For co-imaging of paxillin, RIM and actin, first sequential DNA-PAINT was performed with Imager strands I2-560 and I1-650 (Ultivue) to image RIM and paxillin, respectively, with 100 ms exposure time and subsequently LifeAct-mNeonGreen was added to image actin similar to the imaging of actin and insulin described above. Images were reconstructed using “Detection of Molecules” ImageJ plugin v.1.2.2 (https://github.com/ekatrukha/DoM_Utrecht) (Chazeau et al., 2016).

### Image analysis

#### Colocalization analysis

Pearson’s correlation coefficients were calculated using the Coloc2 plug-in with TIRFM images acquired as described above. A region of interest was picked in both image channels and Pearson’s correlation coefficient was calculated without using a threshold. To rule out non-specific colocalization, the same analysis was done but with one image channel rotated 90 degrees clock-wise.

#### Quantification of focal adhesion complexes

Focal adhesion size was quantified using pFAK TIRFM images acquired as described above. Focal adhesions were detected using “Automated Moments” thresholding followed by particle detection using particle analysis with a minimal size cut-off of 0.15 µm^2^.

Enrichment of LL5β around focal adhesions was quantified using pFAK and LL5β TIRFM images acquired as described above. Focal adhesions were detected using “Automated Moments” thresholding followed by particle detection using particle analysis with a minimal size cut-off of 0.15 µm^2^. The total fluorescence intensity of LL5β was measured in 3 µm-broad areas surrounding each focal adhesion and normalized for surface area. To measure total fluorescence intensity of LL5β outside focal adhesion areas, the opposite of focal adhesion areas was selected by inverting the enlarged focal adhesion areas. Total fluorescence intensity of LL5β outside focal adhesion areas was measured and normalized for surface area. All regions of interests were restricted to areas occupied by cells.

Recruitment of LL5β to focal adhesions upon glucose stimulated insulin secretion was quantified using pFAK and LL5β TIRFM images acquired as described above. Focal adhesions were detected using “Automated Moments” thresholding in ImageJ followed by particle detection using particle analysis with a minimal size cut-off of 0.15 µm^2^. The total fluorescence intensity of LL5β was measured in 1 µm-broad areas surrounding each focal adhesion and normalized for surface area. Finally, the total fluorescence intensity of LL5β was measured in the original focal adhesion areas, normalized for surface area and subtracted from normalized total fluorescence intensity of LL5β in enlarged focal adhesion areas. Only data-points exceeding LL5β fluorescent background signal by 1.5 times where used for plots.

#### Quantification of cluster intensity

LL5β, RIM and GFP-ELKS cluster numbers were quantified using the respective TIRFM images acquired as described above. Images were subjected to Gaussian blur with the radius of 1 pixel and “Unsharp” filtering, followed by “Automated Intermodes” (LL5β) or “Automated Moments” (RIM, GFP-ELKS) thresholding. Next, a watershed-based segmentation was applied, and clusters were detected using particle analysis with a minimal size cut-off of 0.10 µm^2^.

Docked insulin vesicles were quantified using insulin TIRFM images acquired as described above. Images were subjected to Gaussian blur with the radius of 1 pixel and “Unsharp” filtering, followed by “Automated Moments” thresholding. Next, a watershed-based segmentation was applied, and vesicles were detected using particle analysis with a minimal size cut-off of 0.01 µm^2^.

RIM and GFP-ELKS clustering were quantified using the respective TIRFM images acquired as described above. Images were subjected to Gaussian blur with the radius of 1 pixel and “Unsharp” filtering, followed by “Automated Moments” thresholding. Next, a Watershed-based segmentation was applied, and puncta were detected using particle analysis with a minimal size cut-off of 0.10 µm^2^. The obtained regions of interest were subjected to “Nearest Neighbor Distance Calculation” ImageJ plugin and distances were binned in 50 nm bins and plotted as frequency distributions.

Insulin vesicle z-distributions were quantified using insulin confocal images acquired with the setup described above. Z-series images were acquired using a 0.5 µm-step confocal based scan. Images were subjected to Gaussian blur with the radius of 1 pixel and “Unsharp” filtering, followed by “Automated Moments” thresholding. Next, a watershed-based segmentation was applied, and vesicles were detected using particle analysis with a minimal size cut-off of 0.12 µm^2^.

#### FRAP analysis

FRAP experiments with GFP-ELKS-expressing pancreatic islets were performed using the TIRFM setup described above using CoolSNAP HQ2 camera. FRAP was measured by first acquiring 5 frames every 1 s, followed by bleaching a 5-µm-diameter circle in GFP-ELKS patches followed by 5 min imaging with a frame interval of 5 s and 15 min imaging with a frame interval of 10 s. FRAP acquisitions were corrected for the sample drift using 2D cross-correlation function of ‘Correlescence’ ImageJ plugin (ver.0.0.4, https://github.com/ekatrukha/Correlescence). Whole, inner and outer ROI intensity-time measurements were performed using custom written ImageJ macro and averaged in MATLAB. The recovery curves of intensity were background subtracted, corrected for the bleaching caused by the imaging and normalized according to (Phair et al., 2004). To choose the best fitting to the recovery curves, we performed extra sum-of-squares F test between one-phase and two-phase association exponential recovery equations in GraphPad Prism. In both cases (low and high glucose) two-phase association was the preferred model with P<0.0001. Kymographs of recovery areas were built using KymoResliceWide plugin for ImageJ v.0.3 (https://github.com/ekatrukha/KymoResliceWide).

#### Analysis of trajectories of GFP-ELKS clusters and single molecules

GFP-ELKS clusters and single molecule dynamics were imaged using the TIRFM setup described above using Prime BSI camera. To record movement of the clusters, we performed timelapse acquisition every 5 s with 50 ms exposure for 200 frames. Movies were corrected for sample drift using 2D cross-correlation function of “Correlescence” ImageJ plugin (ver.0.0.4, https://github.com/ekatrukha/Correlescence). Cluster detection and linking of trajectories were performed using “Detection of Molecules” ImageJ plugin (https://github.com/ekatrukha/DoM_Utrecht) with linking radius of 0.325 µm per frame and zero frame gap.

For the single molecule acquisition, we selected previously unexposed area of islets and first performed stream acquisition of 1000 frames with 33 ms exposure with full laser power to ensure complete photobleaching of bright clusters. After a short recovery (10-15s) with the same laser power we performed a second acquisition of 1000 frames with 33 ms exposure. These paired acquisitions were used both to track single molecules and quantify stoichiometry of clusters (see below). To separate fast movements of the single molecules, we applied a temporal median filter with the sliding window of 15 frames (∼0.5 s) to the each pixel of the second acquisition (Hoogendoorn et al., 2014, Masucci et al., 2021). To retrieve slow movements, we subtracted the generated results of temporal median filter from all pixels/frames of the original movie using Image Calculator function of ImageJ. Tracking of single molecules in these generated movies was performed using the “Detection of Molecules” ImageJ plugin. Average MSD curves were generated using “msdanalyzer” Matlab class (Tarantino et al., 2014) and imported to GraphPad Prism for linear fitting. Log-log MSD probability density heatmaps were built using a custom written Matlab script.

#### Analysis of the intensity of GFP-ELKS clusters

To determine the number of GFP-ELKS molecules within clusters, we used the paired set of single molecule acquisitions described in the previous section. Cluster intensity distribution *ρ*_clusters_(*I*) was measured from detections in the first frame of unexposed first acquisition. We used amplitude of fitted 2D Gaussian (with an offset background) *I* as a measure of spot intensity. To estimate intensity distribution of single GFP molecules *ρ*_1_(*I*) we selected intensity in the last detection in the trajectories obtained from the slow component of the second acquisition, before complete photobleaching. Both *ρ*_clusters_(*I*) and *ρ*_1_(*I*) were constructed numerically from the individual spot intensity measurements *I*_1_,*I*_2_,..*I*_M_ in the following order. First, we calculated optimal intensity sampling step (bin size) *δI* according to the Freedman-Diaconis rule:

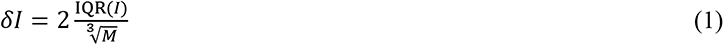

From each fitting of the fluorescent spot, apart from the value of 2D Gaussian amplitude, we determined *σ*, the value of its uncertainty (error). The final *ρ*(*I*) were constructed at intensity steps of *δI* as a sum of normal distributions with means equal to *I*_1_,*I*_2_,..*I*_M_ and standard deviations *σ* _1_*, σ*_2,_…, *σ* _M._
To build corresponding intensity distributions of N colocalized independent GFP molecules *ρ*_N_(*I*), we recursively calculated a series of convolution integrals as described previously (Moertelmaier et al., 2005):

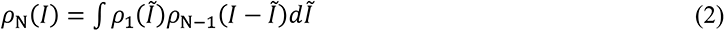

For the numerical implementation, we first approximated discretely sampled *ρ*_1_(*I*) with B-splines. To calculate *ρ*_2_(*I*) we convoluted *ρ*_1_(*I*) with itself using numerical integration with Gauss–Kronrod quadrature formula (Matlab function “quadgk”). After B-spline approximation of *ρ*_2_(*I*), we calculated its convolution with *ρ*_2_(*I*) and so on. Overall, for the purpose of fitting, we constructed GFP N-mers basis consisting of oligomers with N_max_ = 100 GFP molecules. The cluster intensity distribution *ρ*_clusters_(*I*) was fitted with a mixture of oligomers *ρ*_fit_(*I*) using a linear combination of their intensities:

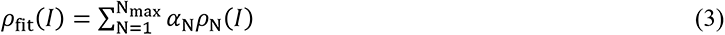

where *α*_N_ stand for the weights of individual distributions, with 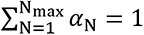 representing fractions of the respective GFP N-mers. The fitting was performed using Matlab “fmincon” function using normalization constraint on the weights. As an optimization function, we used maximum likelihood estimation. Practically, at each fitting step, we minimized a sum of negative log-likelihoods for intensity values used to construct *ρ*_clusters_(*I*) to be sampled from *ρ*_fit_(*I*) at the current minimizing iteration. The procedure was conducted for each field-of-view acquisition and obtained oligomers weight were averaged to produce final histograms.

### Quantification and statistical analysis

All experiments were conducted at least twice. Statistical analysis was performed using GraphPad Prism 9. Statistical details of each experiment, including the statistical tests used, explanation and value of n and precision measures can be found in the figure legends.

## Acknowledgments

We are grateful to M. Jaegle, S. Driegen and B. van der Vaart for technical input in ES cell targeting; J. Kong-a San and A. Maas for blastocyst injection; D. Jaarsma, V. van Dis, E. D. Haasdijk, I. M. Bergen, R. Hendriks, and G. A. Arkesteijn for their help with characterizing *Elks1* knockout mice. We thank N. de Graaf for technical support concerning cell culture and J. Sanes and F. Melchior for the gift of antibodies. The work is supported by the Netherlands Organization for Health Research (ZonMw) TOP grants 91207010 and 91216006, the Netherlands Organization for Scientific Research ALW-VICI grant 865.08.002, Human Frontier Research Grant RGP0001/2016 and European Foundation for the Study of Diabetes (EFSD) grant to A. A.

## Author Contributions

I. N., C. M. vd B. and E. A. K. performed experiments, analyzed data and wrote the paper, F. W. J. B. performed experiments with the human cell line and human pancreatic islets, K. L. Y. generated *Elks1* knockout mice, R. P. T. performed super-resolution microscopy experiments, S. P. organized mouse experiments, B. J. V. performed experiments, L. C. K. set up super-resolution microscopy imaging and data analysis and wrote the paper, F. C. and E. J. P. d K. organized human cell line and pancreatic islet experiments and wrote the paper, E. de G. and C. C. H. organized mouse experiments and wrote the paper, A. A. coordinated the project and wrote the paper.

## Supplemental Movie Legends

**Movie S1. 3D reconstruction of GFP-ELKS in an isolated mouse pancreatic islet.**

3D reconstruction of GFP-ELKS (green) and DNA (blue) in an isolated mouse pancreatic islet imaged by confocal microscopy.

**Movie S2. Dynamics GFP-ELKS clusters in an isolated mouse pancreatic islet.**

An example TIRFM acquisition of GFP-ELKS clusters (top) and the corresponding overlay of trajectories (bottom).

**Movie S3. Dynamics of single GFP-ELKS molecules in an isolated mouse pancreatic islet.**

An example TIRFM acquisition of single GFP-ELKS molecules (top left) and the corresponding overlay of trajectories (bottom left). Right panels illustrate the temporal median filtering method, the splitting of the acquisition into fast (top) and slow (bottom) components.

**Movie S4. Transitions of single GFP-ELKS molecules between diffusive and tethered states.** An example TIRFM acquisition of single GFP-ELKS molecules (left) and the corresponding overlay of trajectories (right).

---

**Figure S1.**
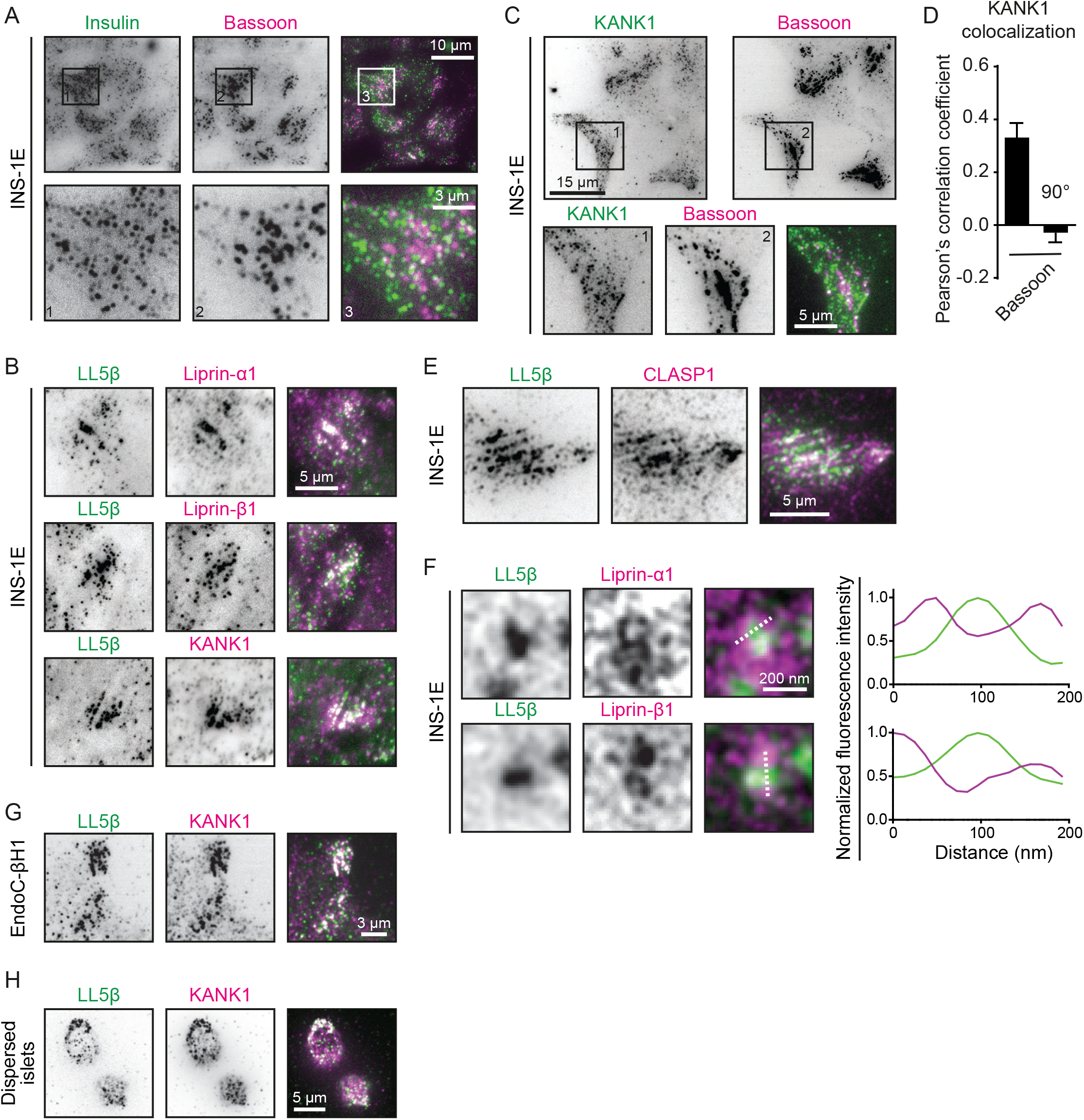
Organization of the insulin secretion sites in INS-1E cells. A. Staining for insulin (green) and Bassoon (magenta) in INS-1E cells imaged with TIRFM. B. Staining for LL5β (green) and liprin-α1, liprin-β1 and KANK1 (magenta) in INS-1E cells imaged with TIRFM. C. Staining for KANK1 (green) and Bassoon (magenta) in INS-1E cells imaged with TIRFM. D. Quantification of colocalization between KANK1 and Bassoon in INS-1E cells. Analysis and display as in Fig. 1B. n=12. E. Staining for LL5β (green) and CLASP1 (magenta) in INS-1E cells imaged with TIRFM. F. Stimulated Emission Depletion (STED) microscopy images of LL5β (green) and liprin-α1 and liprin-β1 (magenta) in INS-1E cells. Intensity profiles along dotted lines are plotted in graphs. G. Staining for LL5β (green) and KANK1 (magenta) in EndoC-βH1 cells imaged with TIRFM. H. Staining for LL5β (green) and KANK1 (magenta) in dispersed human pancreatic islets imaged with TIRFM.

**Figure S2.**
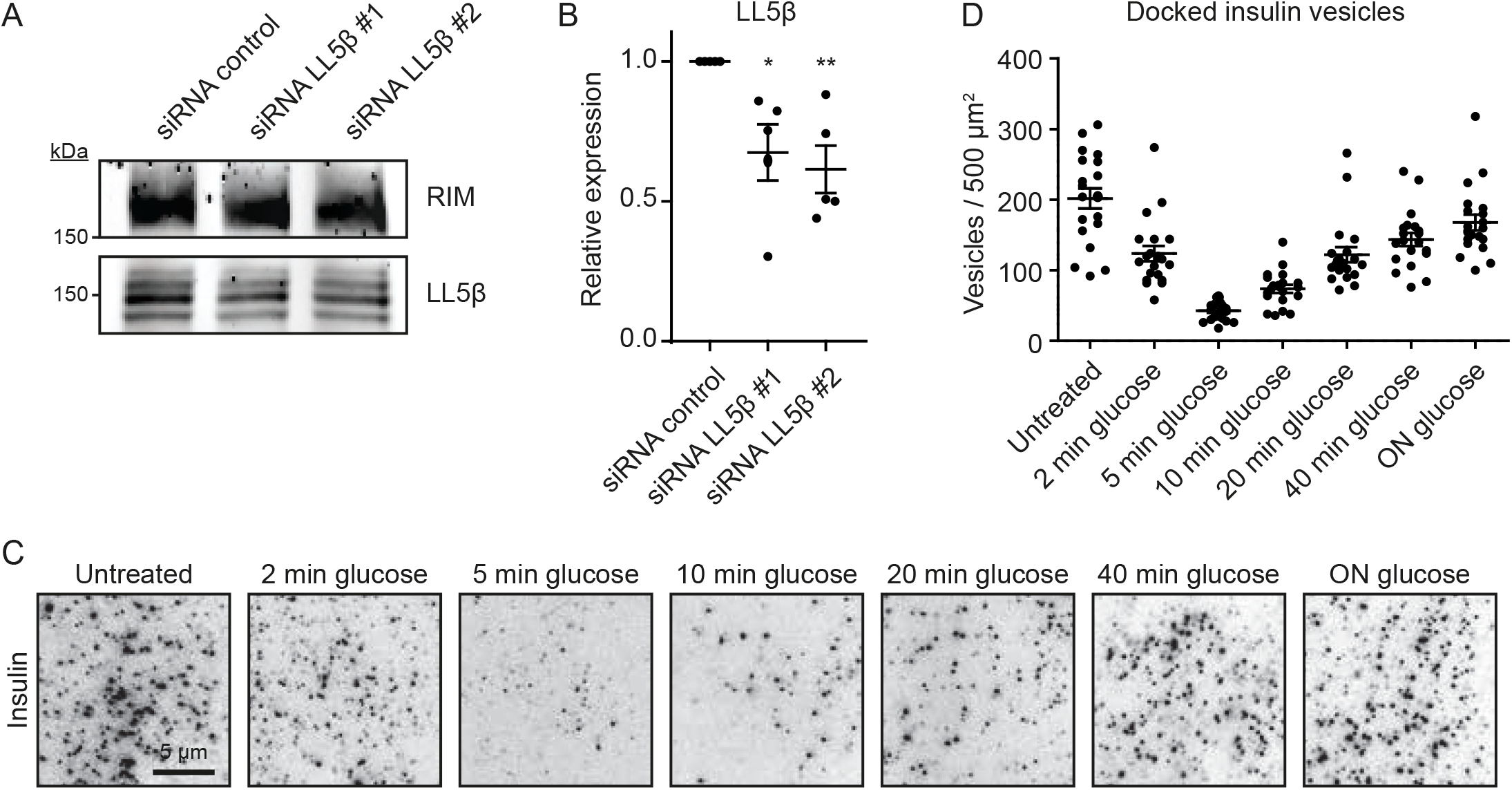
LL5β knock-down and glucose stimulated insulin secretion in INS-1E cells. A. Western blot analysis of RIM and LL5β expression in INS-1E cells treated with siRNAs as indicated. B. Quantification of LL5β expression based on Western blot analysis as shown in (A). *p<0.1; **p<0.01; one-way ANOVA followed by Dunnett’s post-test. Single data points are plotted. Horizontal line, mean; error bars, S.E.M. For all conditions, n=5 C. Staining for insulin in INS-1E cells starved with 2 mM glucose for 4 hours followed by 25 mM glucose stimulation for indicated times and imaged with TIRFM. D. Quantification of docked insulin vesicles in INS-1E cells treated and stained as in (C). Analysis and display as in Fig. 2F. For all conditions, n=20.

**Figure S3.**
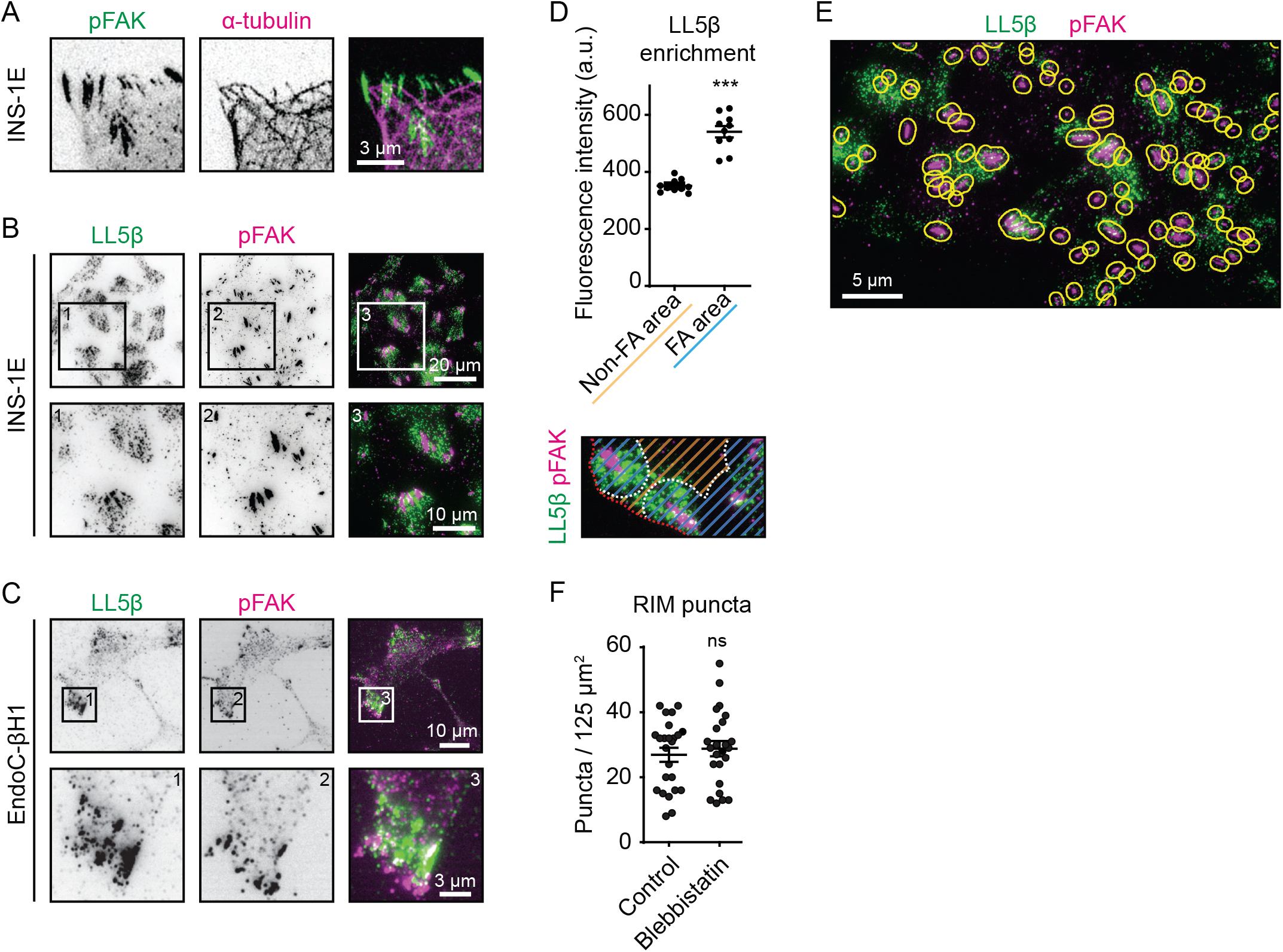
Analysis of the distribution of cortical secretion complexes. A. Stimulated Emission Depletion (STED) microscopy images of phosphorylated FAK (pFAK, green) and α-tubulin (magenta) in INS-1E cells. B. Staining for LL5β (green) and phosphorylated FAK (pFAK, magenta) in INS-1E cells imaged with TIRFM. C. Staining for LL5β (green) and phosphorylated FAK (pFAK, magenta) in EndoC-βH1 cells imaged with TIRFM. D. Quantification of LL5β localization relative to focal adhesions in INS-1E cells. Definition of analyzed cell areas are indicated in scheme. Non-focal adhesion area (orange stripes); focal adhesion area (blue stripes); cell boundary (red dotted line). Single data points are plotted. For both conditions, n=10; ***p<0.001; Mann-Whitney U-test; error bars, S.E.M. E. Analysis example of LL5β (green) localization relative to focal adhesions (pFAK, magenta) in INS-1E cells. Yellow lines indicate areas in which LL5β fluorescence signal was quantified in Fig. 3F. F. Quantification of the numbers of RIM puncta in INS-1E cells treated and stained as in Fig. 3H. Analysis and display as in Fig. 2C. For both conditions, n=24.

**Figure S4.**
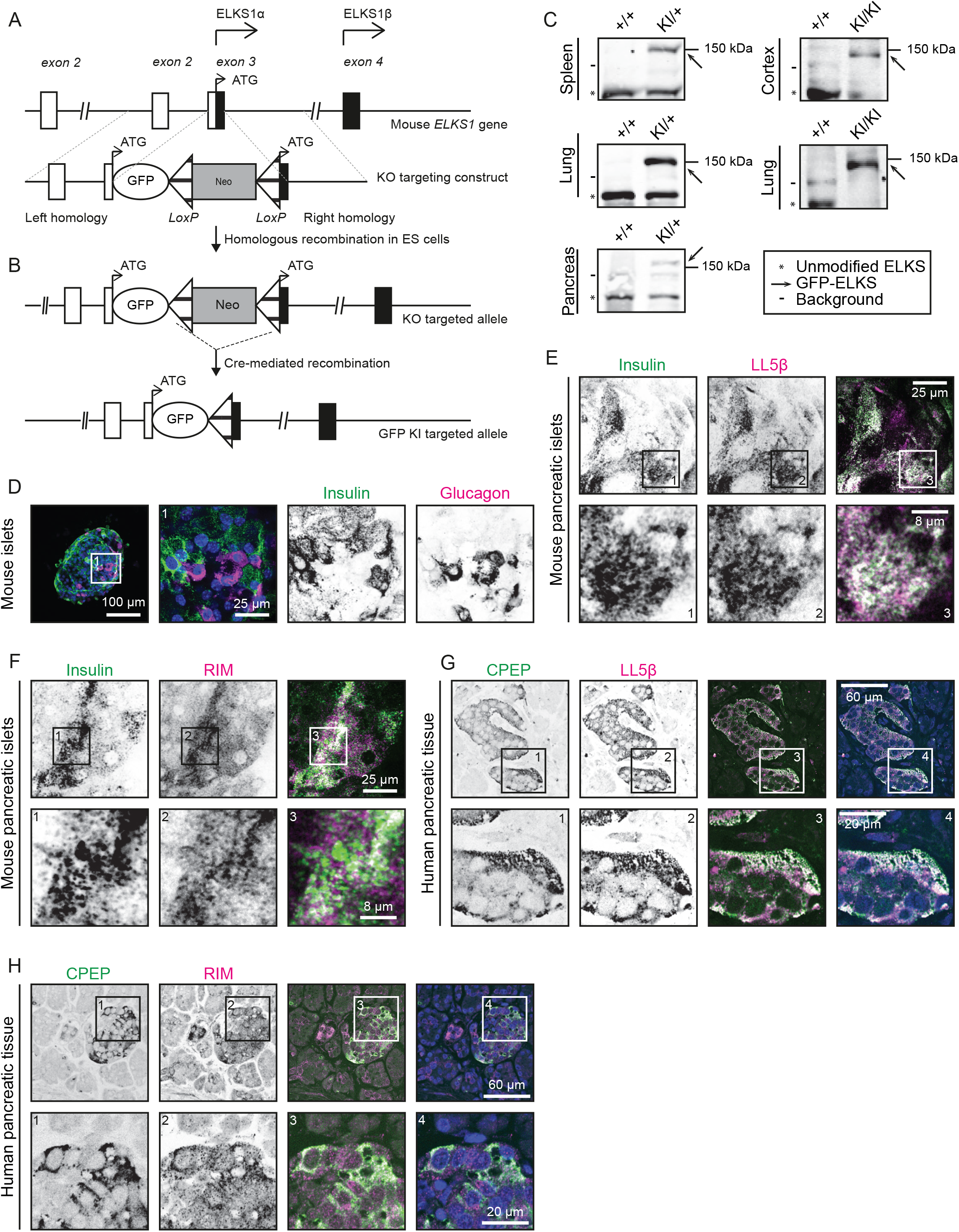
Generation and characterization of GFP-ELKS knock-in mouse line. A. Schematic representation of the ELKS knockout (KO) targeting construct. The top line represents the first four exons of ELKS1 gene on mouse chromosome 6. The bottom line represents the ELKS knockout targeting construct containing GFP, the neomycin resistance cassette (NEO) and two LoxP sites (represented by triangles) flanking both sides of NEO. The KO targeting construct has been inserted into exon 3. B. Schematic representation of the ELKS KO allele and the generation of GFP-ELKS knock-in (KI) targeted allele. The top line shows the ELKS KO targeted allele; after Cre-mediated recombination, the GFP-ELKS KI targeted allele is generated (bottom). C. Western blot analysis of the indicated mouse tissues with ELKS antibodies. The bands corresponding to unmodified ELKS are indicated by asterisks, GFP-ELKS by arrows, and background bands by lines. +/+, wild type; KI/+, heterozygous GFP-ELKS knock-in; KI/KI homozygous GFP-ELKS knock-in. D. Staining for insulin (green), glucagon (magenta) and DNA (blue) in a wild type mouse pancreatic islet imaged by confocal microscopy. E. Staining for insulin (green) and LL5β (magenta) in an adherent region of a mouse pancreatic islet imaged by confocal microscopy. F. Staining for insulin (green) and RIM (magenta) in an adherent region of a mouse pancreatic islet imaged by confocal microscopy. G. Staining for C-peptide (CPEP, green) and LL5β (magenta) and DNA (blue) in human pancreatic tissue imaged by confocal microscopy. H. Staining for C-peptide (CPEP, green) and RIM (magenta) and DNA (blue) in human pancreatic tissue imaged by confocal microscopy.

**Figure S5.**
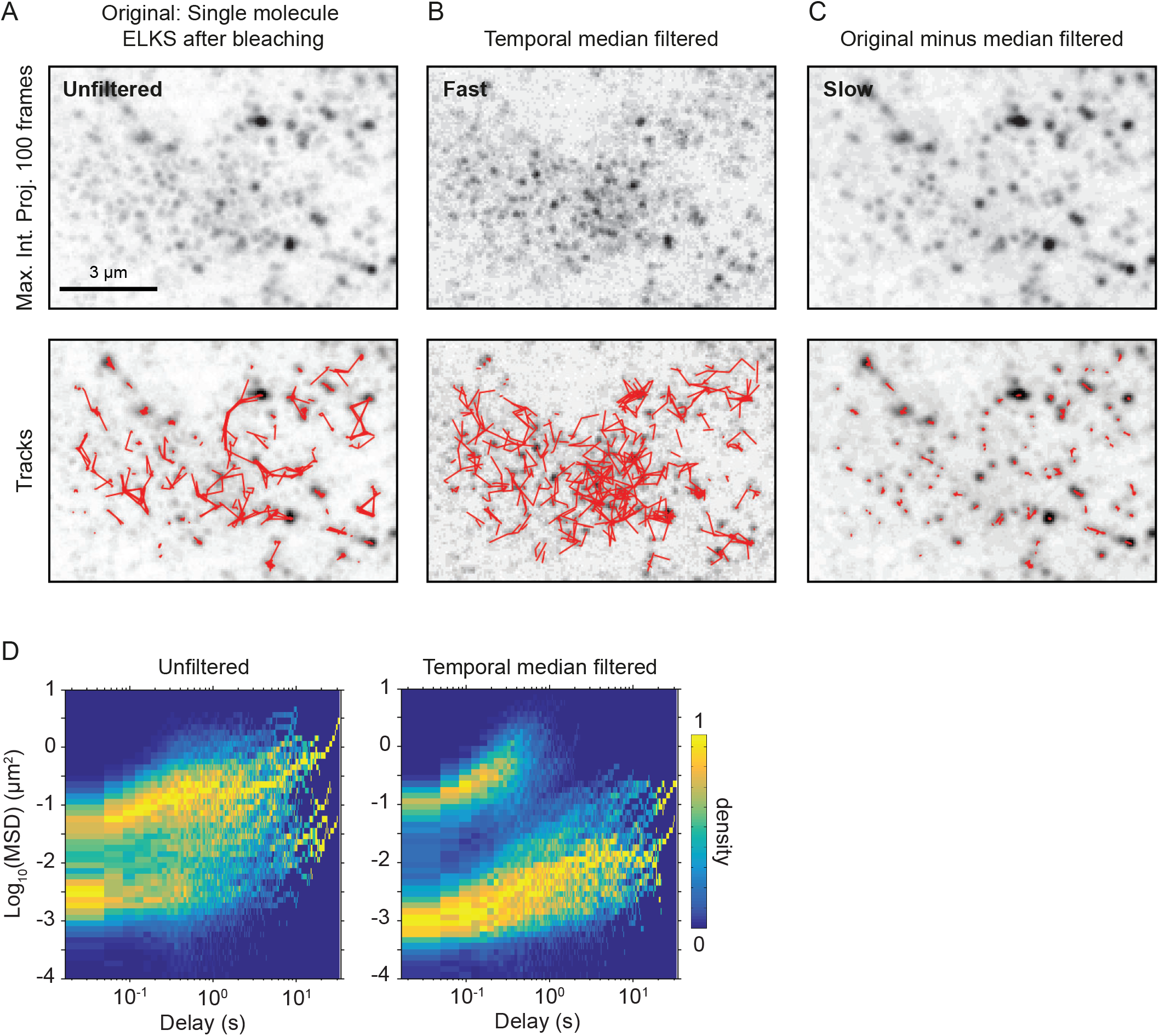
Single molecule analysis of GFP-ELKS in mouse pancreatic islets. A. Representative maximum intensity projection of single GFP-ELKS molecules dynamics (100 frames, 33 ms per frame, top) and the corresponding trajectories (bottom). B. Maximum intensity projection of the movie shown in (A) after application of temporal median filtering with the window size of 15 frames (top) and the corresponding trajectories (bottom). Fast-moving fraction of single molecules is highlighted as a result. C. Maximum intensity projection of the result of frame-by-frame and per pixel subtraction of movie shown in (B) from the movie shown in (A) (top) and corresponding trajectories (bottom). Slow-moving fraction of single molecules is highlighted as a result. D. Heatmap (3D histogram) of MSD values for the trajectories of single GFP-ELKS molecules, tracked with and without temporal median filtering. Histogram values are normalized by the maximum value of each column, corresponding to each time delay bin.

**Supplementary Table 1.** Average FRAP curves fitting parameters

| Fitted value ±<br>error of fit /<br>Condition | Plateau<br>(exchangeable<br>fraction) | Fast halftime<br>(min) | Slow halftime<br>(min) | Percent fast |
| --- | --- | --- | --- | --- |
| Low glucose | $0.42 \pm 0.02$ | $0.93 \pm 0.37$ | $4.8 \pm 1.9$ | $40 \pm 16$ |
| High glucose | $0.65 \pm 0.03$ | $0.67 \pm 0.26$ | $5.5 \pm 1.1$ | $24 \pm 6$ |

## Key Resources Table

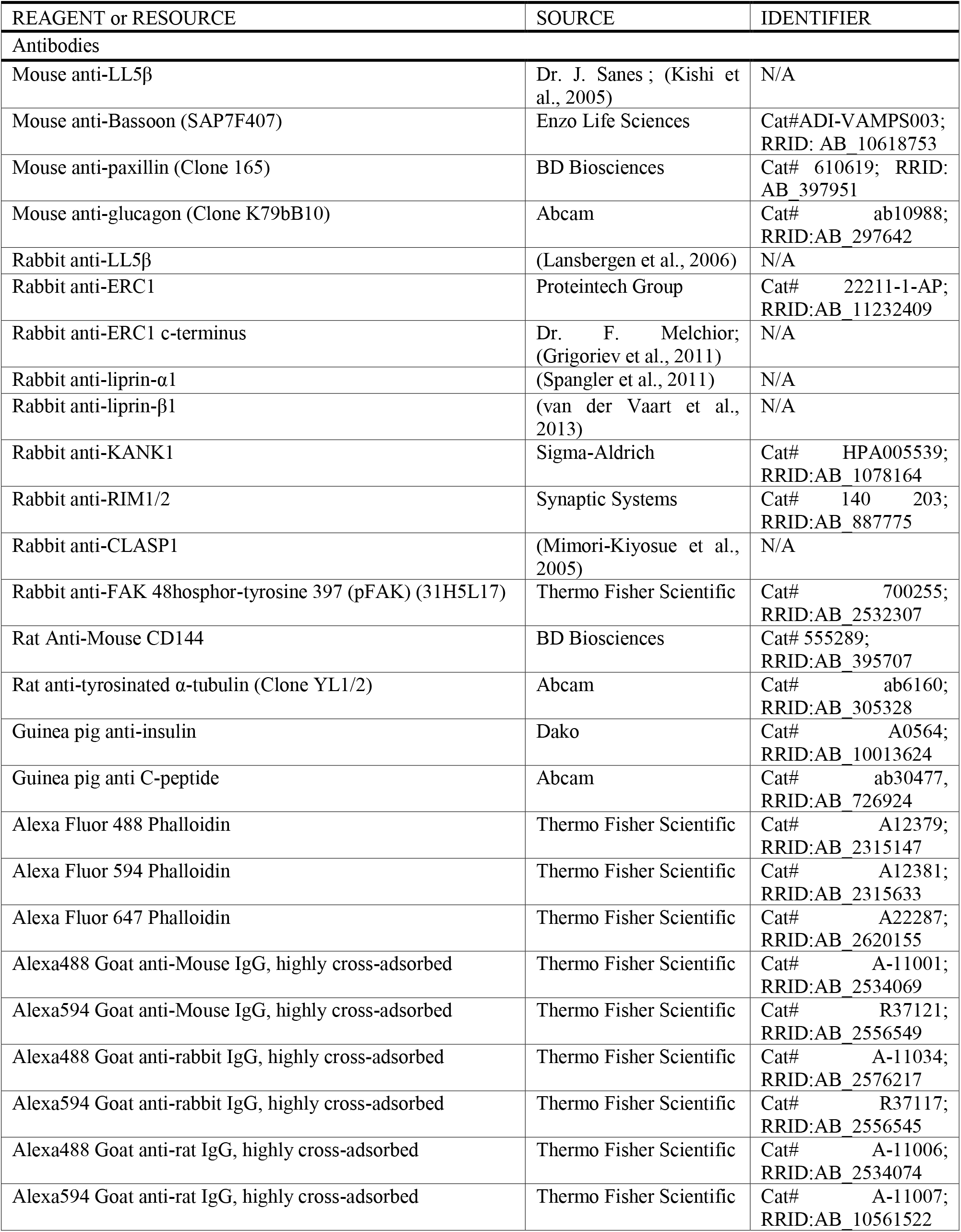
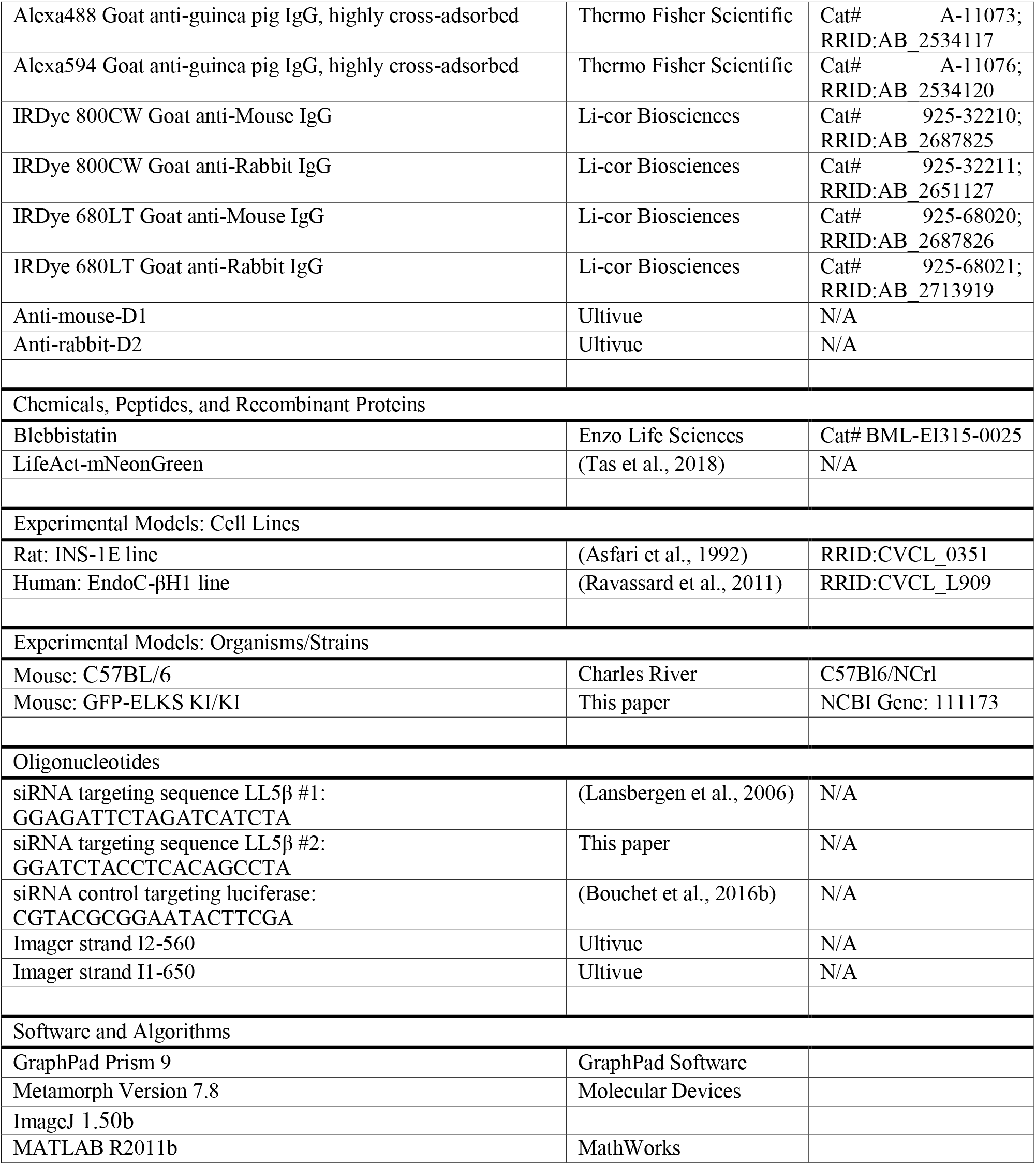

